# Early life stress induces sex-specific changes in behavior and parallel locus coeruleus neuron excitability

**DOI:** 10.1101/2023.12.05.570155

**Authors:** Savannah Brannan, Ben D Richardson

## Abstract

Many psychiatric disorders are associated with specific risk factors, including biological sex, chronic stress, and adversity in childhood, but mechanisms underlying these relationships are unknown. The locus coeruleus (LC) is a brain area that contains adrenergic norepinephrine (NE)-releasing neurons with established sex differences in excitability and stress sensitivity. To understand how adversity in early life affects cognitive/affective behavior and LC physiology, we exposed C57BL/6J mice to a dual-phase (early development and adolescence) early life variable stress (ELVS) paradigm and assessed behavior and LC physiology in early adulthood. ELVS caused females, but not males, to display increased novel environment exploration and reduced preference for sucrose. In addition, ELVS caused elevated activity in a familiar environment, modest deficits in Y-maze performance, and altered attention in an operant task, regardless of sex. A reduction in LC neuron excitability, partly due to an increase in action potential delay time, was also found only in female mice exposed to ELVS, paralleling robust behavioral changes. Pharmacological compensation of these changes in LC activity with reboxetine corrected some ELVS-induced behavioral changes. CRF-induced changes in LC neuron activity were also mediated by different preferential signaling pathways in male and female mice, a potential mechanism for lasting sex-specific changes in LC physiology in response to ELVS. Through this animal model of early life stress, we identified sex differences in behavior and parallel changes in LC neuron excitability and CRF sensitivity, identifying mechanisms involved in determining how stress and sex interact to cause LC activity dysregulation and related behavioral changes.

## Introduction

Adverse childhood experiences (ACEs) have been shown to increase risk for many neurodevelopmental and affective disorders such as attention deficit hyperactivity disorder (ADHD), autism, anxiety and depression. Although mature neurological systems are often resistant to acute stress, prolonged periods of stress in more malleable and immature systems can alter neurodevelopment [1–5]. The more ACEs a child has, the greater the risk for adverse outcomes. Sex is a biological variable involved in ACEs, leading to adverse psychiatric outcomes, with females showing an increased risk for stress-related disorders in comparison to males [6]. Epidemiological studies indicate women may have increased stress-related disorder development risk/symptom exacerbation during life-phases characterized by hormonal changes that include puberty, menses, pregnancy, postpartum, and menopause [2, 3].

Increased ACE exposures leading to attention, memory, affective, and/or mood disorders represent deficits in behavioral domains is shaped by multiple circuits and modulatory neurotransmitters distributed throughout the central nervous system. For several reasons, we hypothesize that the noradrenergic locus coeruleus system (LC) may be a key neuromodulatory brain area responsible for ACE-related changes in behavior. Optimal activity and processing by multiple circuits to shape multiple behaviors depend upon optimal norepinephrine (NE) release from widely-projecting LC_NE_ neurons (e.g. prefrontal cortex, hippocampus, amygdala, etc. [7–15] to modulate attention [16, 17], cognitive flexibility [18, 19], learned behavior [9, 20], anxiety [10, 21] and reward [10, 14], among others [7]. Specifically, work in animal models indicates that LC-NE neuronal projections to the prefrontal cortex (PFC) and hippocampus are involved in maintaining attention, arousal, and aspects of memory formation [11–13, 22, 23] while those to the amygdala (basolateral and central) shape anxiety and emotional learning behaviors [10, 24, 25], and others modulate sensorimotor integration, associative fear learning, response inhibition, and working memory via projections to the cerebellum [26]. Norepinephrine (NE) release is highly dynamic, as appropriate tonic and phasic [16, 17, 27, 28] activity of LC-NE neurons follow an inverted U-shaped function [7, 29, 30]. As such, low and high NE levels both lead to negative outcomes and behavioral changes in cognitive and affective processes.

It is also established that corticotropin-releasing factor (CRF) directly alters the activity of adrenergic neurons in the LC [31–34] and optimal LC neuron activity and related NE release in the prefrontal cortex, amygdala, and hippocampus shape attention, anxiety, and memory [35–39]. Animal model data indicate that, compared to males, female LC neurons have different basal physiological properties [40, 41] and elevated CRF sensitivity that can acutely increase basal NE release from LC neurons [42–44]. These types of basal sex differences in CRF-activated signaling may underpin females’ increased stress-related disorder prevalence [45, 46]. Despite the established role of NE in cognitive and affective behaviors and sex differences in LC and NE dynamics, how early stress interacts with the sexually-dimorphic LC to shape behaviors associated with LC function (i.e. affect, cognition, and locomotion) is poorly understood. To identify and understand the mechanistic relationship between sex and affective, cognitive, and locomotive behavioral changes associated with stress, we developed a two-phase model of stress exposure early in life to determine how these behaviors are affected and if they are related to sex-specific LC physiology changes. Our results suggest an interaction between sex and stress in affective and cognitive behaviors that involve the LC in C57bl6/J (B6J) mice.

## Materials and Methods

### Animals

Male and female C57bl6/J (Jackson Labs, #000664) or Dbh-tdTomato mice (described below) were purchased from Jackson Laboratory and/or bred in-house. To generate mice expressing tdTomato in locus coeruleus noradrenergic neurons under control of dopamine β-hydroxylase (Dbh-tdTomato mice), Dbh-Cre knock-in mice (B6.Cg-*Dbh^tm3.2(cre)Pjen^/J*; stock #033951) provided by Patrician Jensen, NIEHS [47] were crossed with homozygous Ai14 tdTomato reporter mice (B6.Cg-*Gt(ROSA)26Sor^tm14(CAG-tdTomato)Hze^*/J, Stock #007914) provided by Hongkui Zeng, Allen Institute for Brain Science [48]. All mice were group housed under a reversed 12/12-hour light-dark cycle (lights on from 21:00 to 9:00) with *ad libitum* access to food and water. All manipulations and tests were performed during the animals’ active dark phase (9:00 to 21:00), unless otherwise stated. All procedures involving animals were conducted in accordance with protocols approved by the Institutional Animal Care and Use Committee (#2022-100) of Southern Illinois University - School of Medicine.

### Early Life Variable Stress (ELVS) Animal Model

Male and female C57bl6/J mice were subject to either a two-phase (postnatal and adolescent) early life variable stress (ELVS) paradigm or treated as controls with mice in a litter either exposed to ELVS or treated as controls. The ELVS paradigm consisted of two phases of manipulations, first from postnatal day 6-17 and the second from postnatal day 28-42. First, mice in the ELVS group were isolated together from dams for 4hrs/day during postnatal day 6-10 (P6-10) with access to a warming pad, and then weaned early at postnatal day 17 (P17). Pups were supplemented with nutrient gel (Nutra-gel, Bio-Serv, Flemington, NJ, USA) from P17 to P21. ELVS mice were exposed to sequential variable stressors from P28-42, with each stressor conducted twice on their designated day starting with tape restraint (20 min), forced ice-cold swim in a 40 cm x 10 cm filled container (20 min), 2,3,5-Trimethyl-3-thiazoline (TMT) synthetic fox odor exposure (20 min), light cycle disruption by omission of one night of dark phase, and 48 hours of social isolation. Control male and female C57bl6/J mice were not separated from dams and weaned at P21 and were briefly handled 5 times per week for 10-15 minutes beginning at the same time as the stressors (P28) for the ELVS group.

### Behavioral Tests

Behavioral tests were done during the animals’ active dark phase in a dimly lit room (∼15-20 lux) for testing. Mice were habituated to the testing room for 30 minutes prior to completion of behavioral testing. Mouse performance on all behavioral assays, except for the rotarod, were digitally recorded for analysis. Each apparatus was cleaned thoroughly with scent-free disinfectant between animals.

### Open Field Test

Mice were placed in a white acrylic chamber (40 cm x 40 cm wide x 30 cm high square open top chamber) for 30 min with total distance, cumulative time in and number of entrances into center (20 cm x 20 cm) area relative to edges (10 cm border) were analyzed using the EthoVision XT 17 software (Noldus, Wageningen, the Netherlands).

### Y-Maze

Mice were placed in a 3-armed “Y” shaped apparatus with symmetrical arms 35 cm in length, 30 cm wall height and 10 cm lane widths at 120-degree angles from one another. Total alternations, or the number of times the mouse enters 3 arms, were recorded and compared to correct (sequential) alternations, where every arm entered within the 3 entrances had not been entered in the immediate previous entrance.

### Elevated Zero Maze

Mice were placed in the open arm of an elevated circular platform (50 cm diameter with 5 cm wide track) suspended 61 cm from the floor, with two opposing walled (20 cm high) segments and two open segments in between. The amount of time spent in and entries into the open versus closed segments were calculated using Ethovision XT 17.

### Novel Object Recognition and Novel Object Placement Testing

The novel object recognition (NOR) assay was used to assess spatial memory. Following exposure to the open field for 30 min as the chamber acclimation period, the NOR task was performed in the same open field chamber with two phases, each with a 10-minute duration. The first phase was the object habituation period, wherein two identical objects were placed equidistant apart on opposite sides of the chamber. In the second phase, one familiar object from the previous phase was preplaced with a novel object at the same location. Each phase was done on separate days, with the novel object phase occurring 24 h after the same object phase. More time spent with the novel object, was interpreted as increased recall of the familiar object and related to memory and novelty seeking. The amount of time spent with the novel or moved object versus the familiar object is calculated based on the time each mouse spent within 2 cm of the object using Ethovision XT 17. Because this assay was performed in the same arena used as the open field assay, the total distance covered during the second phase (novel object phase) of the assay was also used to determine exploratory activity in a familiar environment to which the mouse had been exposed two previous times.

### Sucrose Preference Test

Two weeks before sucrose preference testing (SPT), mice were progressively water-restricted, first to 4 hours of *ad libitum* water access per day for 2 days and then for 2 hours of access per day for 2 days, followed by two days of 24 hr access. This was repeated twice before limiting water access for 22 hrs prior to sucrose preference testing. For SPT, mice were placed in a clean empty cage for one hour with 2 bottles, one bottle containing 1% sucrose solution and a bottle containing water. The sucrose preference index was determined by comparing the amount of sucrose solution versus water consumed by each individual mouse.

### Rotarod

Motor and vestibular function was evaluated using the accelerating rotarod (3.17 cm diameter, IITC life Sciences, Inc., Woodland Hills, CA, USA). This rod accelerated in rotations per minute (RPM) from 4 10 40 RPM during the span of 5 minutes. Each mouse received 3 trial sessions per day for two consecutive days for a total of 6 trials. A 10 min rest period was given to the mouse between each trial. The time the mouse fell from the rotating rod onto the platform below was recorded. If the mouse spent a reduced fall time latency, it indicated disrupted motor or vestibular function.

### Operant Five-Choice Serial Reaction Time Task (5CSRTT)

For 5SCRTT of attention and learning behaviors, mice were trained using standard 5CSRTT protocols (Asinof and Paine, 2014; Humby et al., 2005) and the apparatus was controlled using ABET software (Lafayette Instruments). Two weeks before training, mice will be water restricted starting at 4 hours of ad libitum access per day, reducing to 2 hours per day. During training sessions, the mice were given sweetened condensed milk as a reward for correctly performing the task and weighed to ensure they were at the proper weight after water restriction. The operant conditioning chamber (30.5 x 24.1 x 29.2 cm) consisted of 2 Plexiglas sidewalls, an aluminum front wall and back wall, and a stainless-steel grid floor will be used to carry out this experiment (Lafayette Instruments). The front wall contained five nose poke apertures (2.5 x 2.2 x 2.2 cm each) with each aperture containing a light-emitting diode (LED) and an infrared sensor cable to detect mouse nose insertions. The mice were trained by progressively shortening the intertrial interval and stimulus duration for 30 minutes daily for 5-7 days per week to reach baseline performance with intertrial interval (ITI) of 5sec and the stimulus duration of 0.8sec. Once mice reached the criterion threshold of 80% correct trials with <30% of trials omitted, they were assessed again on ten different variations with either shorter or longer ITI, shorter or longer stimulus duration, aperture light brightness, sound and light distractor manipulations, and combinations of these changes relative to the standard variables used during training. Response accuracy (% correct) and response omissions (# of premature responses during ITI) for each variable condition, along with the number of premature responses during specific training stages, were evaluated for differences based on mouse sex and treatment as control or exposure to ELVS.

### Reboxetine Administration

In a separate cohort of 7-9-week-old control and ELVS mice, 2 mg/kg reboxetine diluted into apple juice or apple juice alone (100 µl) was given to control and ELVS C57bl6/J mice one hour prior to behavioral testing on SPT, open field, and y-maze. To determine reboxetine effects on these behaviors, one behavior was evaluated each day on consecutive days with pre-test apple juice/reboxetine administered each day 1 hr prior to each assay. The order of the behavioral assays was kept the same for all mice.

### Ex vivo Acute Brian Slice Preparation

Animals were anesthetized with 4% isoflurane, followed by intracardial perfusion with ice-cold oxygenated N-Methyl-D-glucamine (NMDG) artificial cerebrospinal fluid (NMDG-ACSF), which contained (in mM): 92 NMDG, 2.5 KCl, 0.5 CaCl_2_, 10 MgCl_2_, 1.2 NaH_2_PO_4_, 30 NaHCO_3_, 20 HEPES, 25 D-glucose, 2 ascorbic acid, 2 thiourea, and 3 sodium pyruvate and had an osmolarity of 300–310 mOsm with pH adjusted to 7.3–7.4 with HCl. Coronal slices (250 μm thick) that included the LC were sectioned in ice-cold NMDG-ACSF using a Compresstome vibrating microtome (Processionary Instruments, LLC) and were transferred to a holding chamber containing NMDG-ACSF at 35 °C where the NaCl concentration increased steadily over 25 min. After 30 min, slices were transferred to a modified HEPES-based ACSF (HEPES-ACSF, 35 °C), which contained (in mM): 92 NaCl, 2.5 KCl, 2 CaCl_2_, 2 MgCl_2_, 1.2 NaH_2_PO_4_, 30 NaHCO_3_, 20 HEPES, 25 D-glucose, 2 ascorbic acid, 2 thiourea, 3 sodium pyruvate, and 3 myo-inositol and had an osmolarity of 300–310 mOsm with pH 7.3–7.4. After incubating slices in HEPES-ACSF at 35 °C for 1 hr, slices were maintained in HEPES-ACSF at room temperature until being transferred to the recording chamber where they were continuously perfused at a 3-5 ml/min with oxygenated ACSF, which contained (in mM): 125 NaCl, 2.5 KCl, 2 CaCl_2_, 2 MgCl_2_, 1NaH_2_PO_4_, 26 NaHCO_3_, 20 D-glucose, 2 ascorbic acid, and 3 myo-inositol and had an osmolarity of 310–320 mOsm with pH 7.3–7.4.

### Ex vivo Acute Brain Slice Whole-Cell Electrophysiology

Neurons in the LC were visualized using an upright microscope (Olympus BX51WI) with a 40x water-immersion objective. The microscope was able to detect interference contrast and imaged via an infrared-sensitive camera connected to a video monitor. Whole-cell patch-clamp recordings were performed at 32 °C maintained with an in-line solution heater from visually identified LC neurons using the 4^th^ ventricle as a landmark. Noradrenergic neurons in the LC were identified by the presence of tdTomato in Dbh-tdTomato mice and action potential properties of putative noradrenergic LC neurons in C57bl6/J mice were verified to be similar to Dbh-tdTomato+ neurons.

Using a horizontal puller (P1000, Sutter Instruments, Novato, CA, USA), patch pipettes were pulled from filamented borosilicate glass capillaries (outer diameter 1.5 mm, inner diameter 0.86, Sutter Instruments), having a tip resistance of 5–7 MΩ when filled with potassium gluconate-based internal solution that contained (in mM): 139.6 mM K-gluconate, 0.4 mM KCl, 4 mM NaCl, 0.5 mM CaCl_2_, 10 mM Hepes, 5 mM EGTA free acid, 4 mM Mg-ATP, 0.5 mM Na-GTP with an osmolality 285–290 mOsm and the pH adjusted to 7.2-7.3 with KOH (E_Cl_ = -85 mV). The calculated liquid junction potential of 15.5 mV was left uncorrected. For experiments assessing the role of ATP-sensitive potassium channels, all nucleotide salts were removed from the internal solution. For voltage-clamp recordings of synaptic activity, 5 mM QX-314 was included in the internal solution (iso-osmotically replaced NaCl) to block voltage-gated Na^+^ channels. All signals were acquired at 20 kHz and low-pass filtered at 10 kHz via a MultiClamp 700B amplifier. Data were collected from the neurons with an input resistance >100 MΩ. If the series resistance for a given recording was >30 MΩ or changed by more than 20%, the data were rejected for analysis. The current and voltage signals were recorded and digitized with a MultiClamp 700B amplifier and Digitdata 1440 controlled by Clampex 10 data acquisition software (Molecular Devices, San Jose, CA, USA). For 5-6 weeks old LC excitability and CRF response recordings, baseline excitability was collected 5 minutes after whole-cell and after evoked current injections (from -30 pA, increasing by 10 pA up to 100 pA with 10 seconds between each sweep) to ensure stable recordings and complete dialysis of the cytosolic space with intracellular solution. CRF responses were collected within 10 minutes following 500 nM CRF (human, rat, AnaSpec, Fremont, CA, USA) applied in the ACSF bath application. For recordings from 7-8-week old mice, spontaneous firing rates were collected first during a 3-5-minute free run, with sweeps excluded if they were 20% different from the average firing rate. Evoked firing responses to current injection steps were next performed using the same protocol as previously described. IN current-clamp mode, the neuronal membrane potential was kept at -60 mV by injecting an additional holding current, then the same current injection protocol was performed. The action potential threshold was determined by taking the average potential at which the dV/dt reached 20mV/ms when neurons fired spontaneously. To determine the resting membrane, an all-points histogram was generated for the spontaneous firing recording and the peak of the histogram (0.5 mV bins) was identified as the resting potential in highly active neurons. Neuronal membrane resistance was determined based on the holding current required for 5 mV step in voltage-clamp mode.

### Quantification and statistical analysis

All data were evaluated with two- or three-way ANOVAs or repeated measures ANOVAs (RMANOVAs), results of which are reported in **Supplementary Table 1**. If Spearman’s test for heteroscedasticity and the Shapiro-Wilk test for normality were not significant (*p* > 0.05), paired t-tests (drug effects), unpaired t-tests (stress effects), or Fisher’s Exact tests were used for pairwise comparisons as indicated. If either Spearman’s or the Shapiro-Wilk test were significant (*p* < 0.05), a Wilcoxon Signed Rank Test for paired data (dug effects) or Mann-Whitney U test for unpaired data were used for pairwise comparisons. Alpha levels of *p* < 0.05 were considered significant for all main effects and pairwise comparisons, and *p* < 0.10 was considered significant only for interactions when the main effects of stress had *p* < 0.05 in ANOVAs (**Figure 1c** and **2f**). All results are expressed as the mean ± SEM and individual data points represent individual animals in behavioral assays or neurons in cellular assays. Analyses and graphing were conducted in Clampfit 10.0 (Molecular Devices, San Jose, CA, USA), Easy Electrophysiology (Easy Electrophysiology Ltd, London, UK), Prism (10.0.2, Graphpad, Boston, MA, USA), and Igor Pro 8 (WaveMetrics, Lake Oswego, OR, USA)

**Figure 1.**
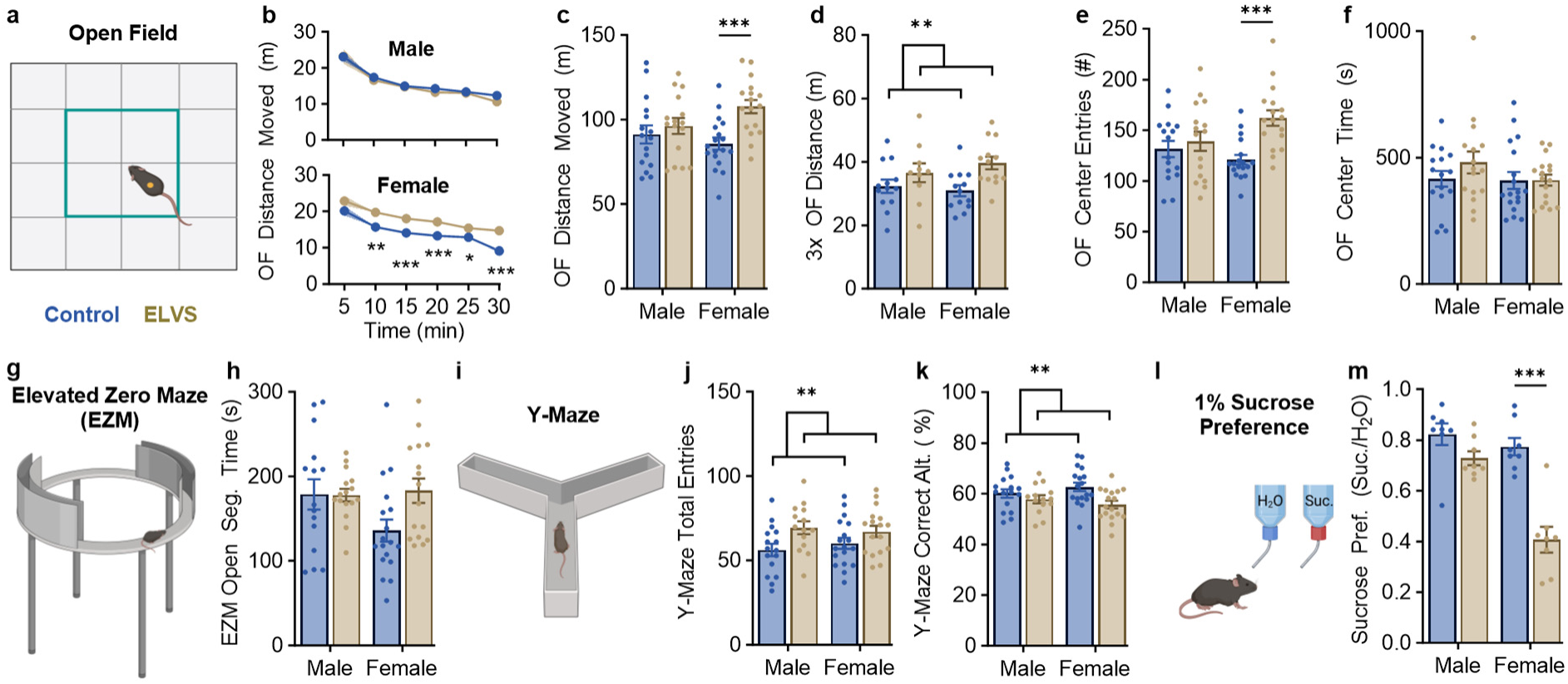
ELVS causes sex-independent and female-specific changes in cognitive and affective behavior. (**a-c**) Total distance (meters, m) moved in the open field (OF) assay (**a**) across the entire 30 min session (**b**; females: U = 48.0 – 92.0, *p* = 0.00030 – 0.045) and in total (**c**; male: *t*(30) = 0.70, *p* = 0.49; female: *t*(33) = 4.08, *p* = 0.0003). (**d**) Total distance (m) moved in OF apparatus during the third exposure (3x OF) for 10 minutes (combined: *t*(47) = 3.13, *p* = 0.0030). (**e**, **f**) The number of entries into (**e**; male: *t*(30) = 0.63, *p* = 0.53; female: *t*(33) = 4.63, *p* < 0.0001) and total time spent (**f**) in the center of the OF arena. (**g**, **h**) Time (seconds) spent (**h**) in the elevated zero maze (EZM, **g**) open arm. (**i**-**k**) Y-Maze (**i**) total number of arm entries (**j**; combined: *t*(63) = 2.83, *p* = 0.0063) and correct arm alternation percentages (**k**; combined: *t*(63) = 2.94, *p* = 0.0046). (**l**, **m**) Sucrose preference (**l**) index values as the ratio of 1% sucrose/H_2_O only (**m**: male: *t*(14) = 1.89, *p* = 0.08; female: *t*(14) = 5.99, *p <* 0.0001). All results are presented as mean ± SEM for each group with circles representing data from individual mice. **a**-**k**: N = 10-18 mice (6 weeks) per group. **m**: N = 8 mice (9 weeks) per group. \**p* < 0.05, \*\**p* < 0.01, \*\*\**p* < 0.001 with two-tailed unpaired t-test or Mann-Whitney U test for pairwise comparisons.

**Figure 2.**
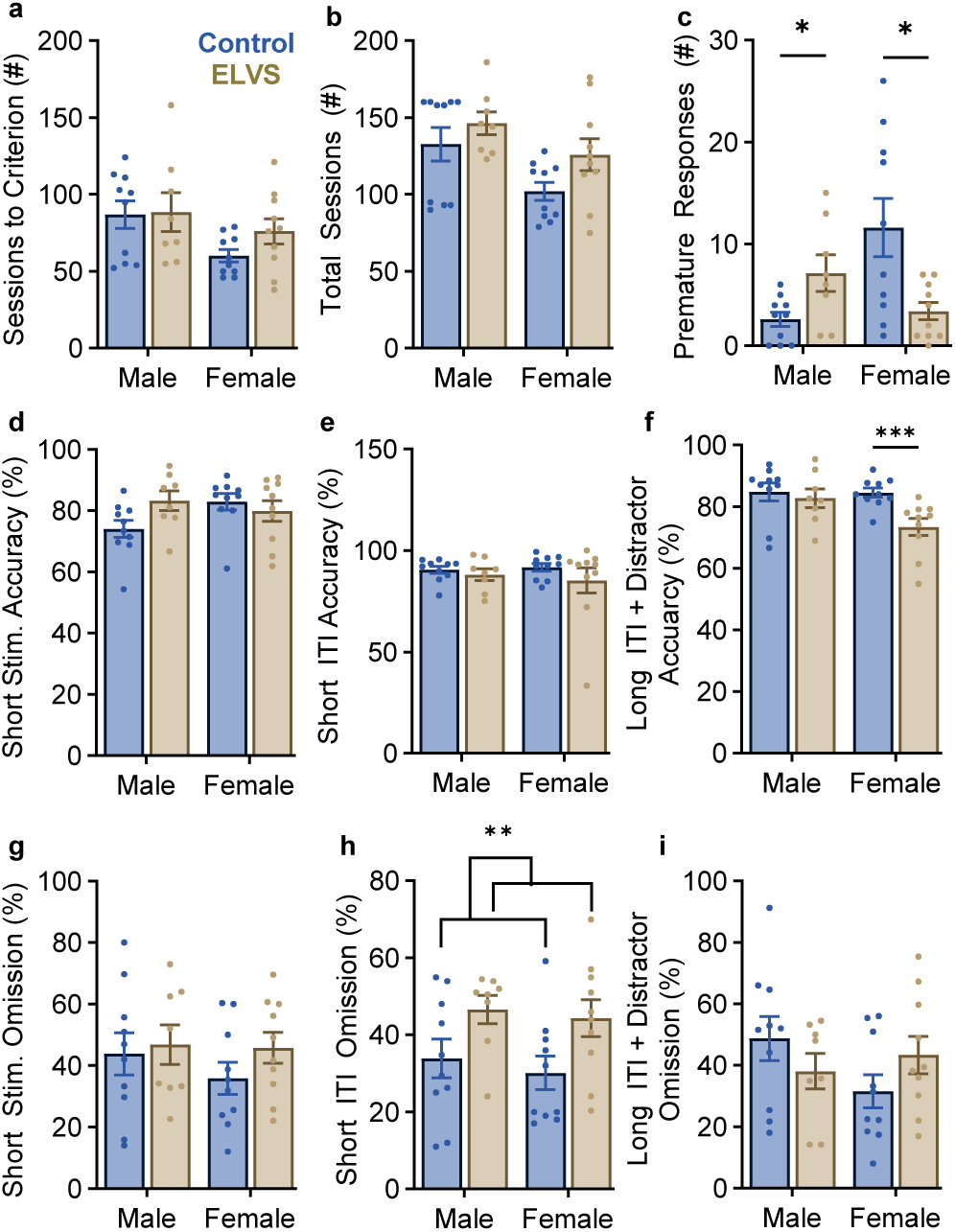
ELVS leads to sexually dimorphic changes in impulsivity and female-specific distractibility in 5-choice serial reaction time task (5CSRTT). (**a**) Days needed to meet criteria during 5CSRTT training. (**b**) Days required to complete training and all session variation tests (male: *t*(16) = 0.98, *p* = 0.34; female: *t*(18) = 2.02, *p* 0.059). (**c**) Number of premature responses during the first training transition session in 5CSRTT training with reduced stimulus duration and intertrial interval (males: U = 15.0, *p* = 0.024; females: U = 22.0, p=0.033). Trial accuracy % (**d**-**f**) and omission % (**g**-**i**) in test sessions with shortened stimulus duration in **d** and **g**, shortened intertrial interval in **e** and **h** (combined: *t*(36) = 2.97, *p* = 0.0053), and lengthened intertrial interval with a simultaneous sound distractor in **f** (male: U = 30.5, *p* = 0.42; female: U = 9.0, p=0.0010) and **i**.

## Results

### ELVS induces hyperactivity in a non-novel environment and impaired short-term memory with female-specific increased exploratory and anhedonia-like behavior

To determine how early life stress impacts affective, cognitive, and locomotive behaviors of male and female mice, we exposed C57Bl/6J (B6J) mice to a two-phase stress paradigm during early life (P6-16) and adolescence (P28-P42) as described. At 6-7 weeks of age, male and female control (handled daily) and ELVS mice were evaluated on a behavioral battery, including open field (OF), Y-maze, elevated zero maze (EZM), novel object recognition (NOR), sucrose preference (SP), and rotarod. Female ELVS mice displayed an increase in OF distance moved (**Figure 1a-c**) when compared to female controls that persisted across the 30 min trial and was absent in male mice. In contrast, total locomotion in the familiar open field after the third exposure (habituation phase of NOR) was significantly increased in ELVS-exposed mice, regardless of sex, indicating the development of hyperactivity in a familiar environment due to ELVS exposure. Female ELVS mice also entered the center of the open field arena more frequently than control females (**Figure 1e**), consistent with total OF locomotion. Mice of both sexes spent similar amounts of time in the center of the OF, regardless of ELVS exposure (**Figure 1f**), suggesting a lack of increased anxiety-like behavior despite ELVS exposure. Consistent with OF center time and absence of anxiety-like behavior, there were no significant differences between groups in the cumulative duration of time spent in the open arms of the elevated zero maze (**Figure 1g, h**). In the Y-maze (**Figure 1i-k**), ELVS-exposed mice of both sexes demonstrated increased arm entries (**Figure 1j**) and decreased correct arm alternation sequences (**Figure 1k**), indicating modestly impaired short-term memory relative to control mice. There were no significant sex- or stress-related differences observed in 24 hr novel object recognition (NOR, **Supp.** Figure 1a-d) or rotarod latency to fall times (**Supp.** Figure 1e-g), which tested long-term (24 hr) memory and motor function/coordination, respectively.

Following innate behavioral testing at 6 weeks of age, 7-week-old mice were water restricted for two weeks, then completed sucrose preference testing. Only female ELVS mice displayed a significant reduction in sucrose preference (**Figure 1m**) relative to control mice, which suggests ELVS induces an anhedonia-like state in female mice. Overall, in mice of both sexes at 1.5 – 2 months of age, ELVS exposure led to increased exploratory activity and impaired short-term memory, both ADHD-relevant behavioral domains. In parallel, hyperactivity and anhedonia-like behavior were observed specifically in female mice, indicating changes in affective behavior due to a previous exposure to dual-phase ELVS.

### ELVS exposure results in male-specific increases in impulsivity and female-specific deficits in attention

Mice continued to be water restricted for the five-choice serial reaction time task (5CSRTT) training and testing. There was no effect of stress on the time necessary for mice to reach the criterion threshold (>80% accuracy and <30% omitted trials; **Figure 1a**). Although there were group effects of stress and sex on the total time to complete all training and testing sessions (**Figure 1b**), there were no significant ELVS effects on these metrics for either sex. As a measure of impulsivity, we evaluated the number of premature nosepoke responses for the first training trial when the stimulus duration and intertrial interval were shortened (**Figure 1c**), where we found that premature responses increased in males exposed to ELVS in contrast to being decreased in ELVS-exposed females. Once the mice had been trained to criterion on standard parameters, mice were evaluated in 5CSRTT sessions with varied parameters across ten different test sessions. Accuracy and trial omissions were similar for all groups on most test sessions (**Supp.** Figure 2), including those with short stimulus duration (**Figure 1d**, **g**). However, when the intertrial interval was reduced, ELVS-exposed mice of both sexes performed with similar accuracy (**Figure 2e**), but omitted significantly more trials (**Figure 2h**). 5CSRTT response accuracy was also reduced in ELVS female mice when session parameters were challenging with longer intertrial intervals in the presence of an audio distractor and the target brightness was varied (**Figure 2f**), with no changes in omissions (**Figure 2i**). These behavioral data demonstrate how ELVS exposure leads to male-specific increases in impulsivity while impairing specific attention parameters in ELVS females, suggesting sexually dimorphic effects of ELVS on specific neural circuits responsible for these behavioral changes.

### Corticotropin-releasing factor (CRF) directly excites noradrenergic neurons based on ELVS exposure and via different signaling pathways in male and female mice

Based on previous studies showing the LC’s involvement in affective and attentive behaviors [35, 49, 50] and sex differences in second messenger pathways mediating CRF effects in LC neuron [32], we further probed this sex difference during adolescence to determine the degree to which CRF-evoked changes in LC excitability is dependent on sex and ELVS exposure. To determine the accuracy of targeting putative noradrenergic LC neurons recorded in wildtype B6J mice, we assess CRF (500 nM) effects on LC neuron activity using transgenic mice expressing tdTomato driven by noradrenergic marker, dopamine β-hydroxylase (Dbh-tdTomato mice) (**Figure 3a** & **b**). Baseline and CRF-induced changes in firing behavior of tdTomato+ noradrenergic LC neurons were similar to baseline and CRF-induced changes in B6J mice of both sexes (**Figure 3c-g**). Because the generation of Dbh-tdTomato mice relies on a knock-in of Cre in place of one copy of the *Dbh* gene, mechanistic studies were not performed in these mice due to potential compensatory changes that may occur in the absence of one copy of *Dbh*. In B6J mice, we controlled for whole cell configuration alterations in the internal cell conditions by performing cell-attached recordings of responses to CRF bath applications, with CRF responses similar to whole cell results (**Supp.** Figure 3a). There were also no significant changes in spontaneous firing rate when cells were held in the whole cell condition (**Supp.** Figure 3b), suggesting that this did not confound the observed firing changes upon CRF application.

**Figure 3.**
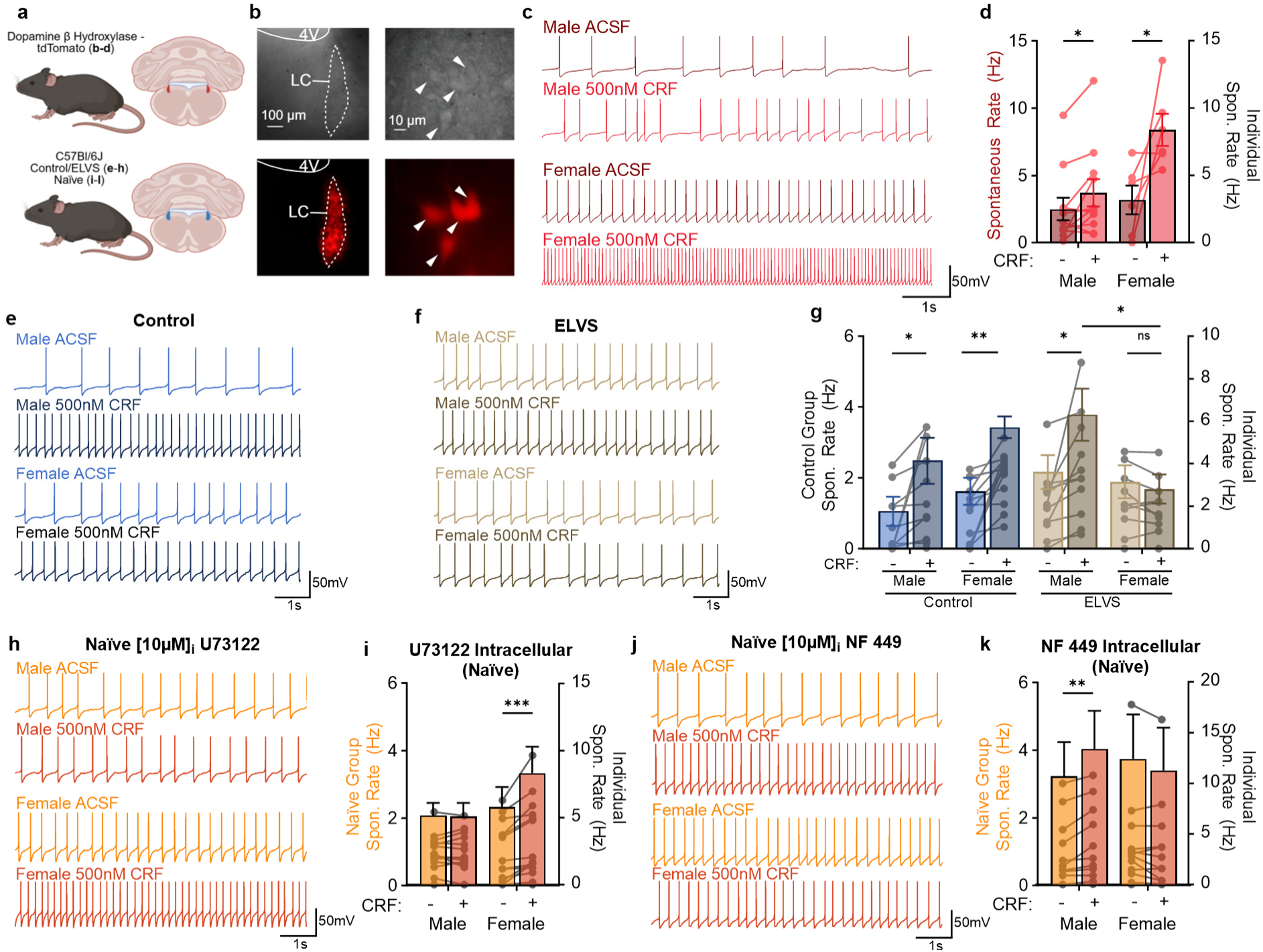
CRF excites LC neurons via different pathways in male and female mice and is lost in female mice after stress exposure. (**a**) Schematic of animals used and brain slice approach for recording form LC neurons in mice expressing tdTomato in dopamine β-hydroxylase (DβH)-expressing neurons, control, ELVS-exposed, or naïve mice. (**b**) DIC (top) and fluorescent tdTomato (bottom) image of the LC below the fourth ventricle (left) and a high magnification image of LC neurons in DβH-tdTomato mice. (**c**, **e**, **f**, **h**, **j**) Representative paired membrane voltage traces of spontaneous action potentials generated by LC neurons in ACSF and after bath application of 500nM CRF from DβH-tdTomato (**c**), control C57 (**e**), and ELVS C57 (**f**) mice, or naïve mice with 10µM U73122 (**h**) or 10µM NF 449 (**j**) in the intracellular solution. (**d**, **g**, **i**, **k**) Group spontaneous action potential frequency of LC neurons in ACSF and after bath application of 500nM CRF from DβH-tdTomato (**d**; male: *t*(10) = 2.55, *p* = 0.029; female: *t*(5) = 2.61; *p* = 0.048), control and ELVS C57 mice (**g**; Control male – CRF: *p* = 0.044; ELVS male – CRF: *p* = 0.0015; Control female – CRF: *p* = 0.020; female control CRF vs. ELVS CRF: *p* = 0.045; ELVS female-CRF: p=0.17), or naïve mice with 10µM U73122 (**i**; male: *W* = 3, *p* = 0.93; female: *W* = - 91, *p* = 0.0002) or 10µM NF 449 (**k**; male: *t*(10) = 3.44, *p* = 0.0063; female: *t*(12) = 1.53; *p* = 0.15) in the intracellular solution. In **d**, **g**, **i**, and **k**, the left y-axis corresponds to mean ± SEM group bars and right y-axis corresponds to individual paired values for each individual neuron. All group data are presented as mean ± SEM (bars) for each group, with connected circles representing paired pre- and post-CRF values from individual neurons. Two-(**d**, **i**, **k**) or three-way (**g**) RMANOVAs were used for group comparisons, followed by a paired t-test, Wilcoxon Signed-Rank test for single pairwise comparisons in two-way RMANOVAs or Bonferroni posthoc in three-way RMANOVAs to assess CRF effects. n = 6-11 neurons from N = 2-4 mice (6-8 weeks old) for **c** and **d,** or n = 11-16 neurons from N = 3 mice (5-7 weeks old) for **e**-k. \**p* < 0.05, \*\**p* < 0.01, \*\*\**p* < 0.001 for pairwise comparisons.

To determine how ELVS exposure impacts CRF-induced LC neuron excitability changes, we performed the same protocol of bath-applying CRF (500 nM) onto LC neurons of ELVS mice at 6 weeks of age after completion of the ELVS protocol. While ELVS male LC neurons were still responsive to CRF, there was a significant interaction between sex and stress on the effects of CRF on spontaneous firing, with ELVS female LC neurons no longer responding by a change in firing rate (**Figure 3e-g**). Sex differences in LC neuron CRF sensitivity may depend on sexually dimorphic variations in CRF actions on LC neurons, which may shift after exposure to stress of different types and timing in early life.

Although it has previously been reported that female LC CRFR1 receptor is predominantly coupled to the G_s_ pathway [51–53], since CRFR1 can couple to G_s_, G_q_, and G_i_ pathways, we went on to determine the role of G_q_ and G_s_ signaling in CRF-induced increases in LC neuron excitability in both sexes. To determine G_q_ pathway dependence of CRF effects on excitability, a protein lipase C (PLC) inhibitor 10 µM U73122 was added to the internal solution. In the presence of the PLC inhibitor, bath application of 500 nM CRF no longer increased spontaneous firing of male noradrenergic neurons (**Figure 3h** & **i**). In contrast, the inclusion of the PLC inhibitor had no effect on CRF-induced changes in the spontaneous firing behavior of female noradrenergic LC neurons (**Figure 3h & i**). 10 µM NF 449, a Gα_s_ inhibitor [54], was next added to the internal pipette solution to determine dependence of G_s_-mediated signaling in CRF-induced excitability changes. CRF no longer significantly increased female LC neurons’ spontaneous firing in the presence of NF 449, but CRF still significantly increased spontaneous firing in LC neurons from males (**Figure 3j** & **k**). CRF did not have a robust effect on evoked firing in any groups or pharmacological treatments, except in naïve males with NF449 added to the internal (**Supp.** Figure 4), which is consistent with changes in spontaneous firing in response to CRF. Together, these data suggest that CRF preferentially signals through G_q_ signaling pathways in males and G_s_ signaling pathways in females, which may explain how stress signaling may lead to sexually dimorphic long-term changes.

### ELVS Female LC Neurons are less excitable after stress and show increased action potential delay attributed to A-type potassium channels

Given the sexual dimorphism of the behavioral phenotype produced by ELVS, sex differences in LC neurons’ CRF sensitivity, female-specific ELVS effects on LC neuronal CRF sensitivity, and the lack of excitatory or inhibitory synaptic activity differences after stress (**Supp.** Figure 5), we next assessed current clamp recordings from putative LC noradrenergic neurons in *ex vivo* acute brain slices to determine whether ELVS produced persistent sex-specific changes in excitability. At 6 weeks of age, immediately following stress, there were no clear changes observed in LC neuron excitability (**Supp.** Figure 6), although at 7-8 weeks of age we found robust ELVS-induced female-specific changes in LC neuron activity (**Figure 4**). Noradrenergic LC neurons from female ELVS mice, but not males, displayed reduced spontaneous firing rates when compared to controls (**Figure 4a** & **b)** with no changes in resting membrane potential (V_m_; **Figure 4c**). Surprisingly, the membrane resistance was higher and the action potential (AP) threshold more depolarized in ELVS female LC neurons (**Figure 4d** & **e**). Both male and female ELVS LC neurons displayed modest, but significantly increased AP half-width (**Figure 4g**). LC neurons from ELVS females also elicited fewer action potentials in response to current injection compared to controls, a change which was absent in males (**Figure 4h-l**).

**Figure 4.**
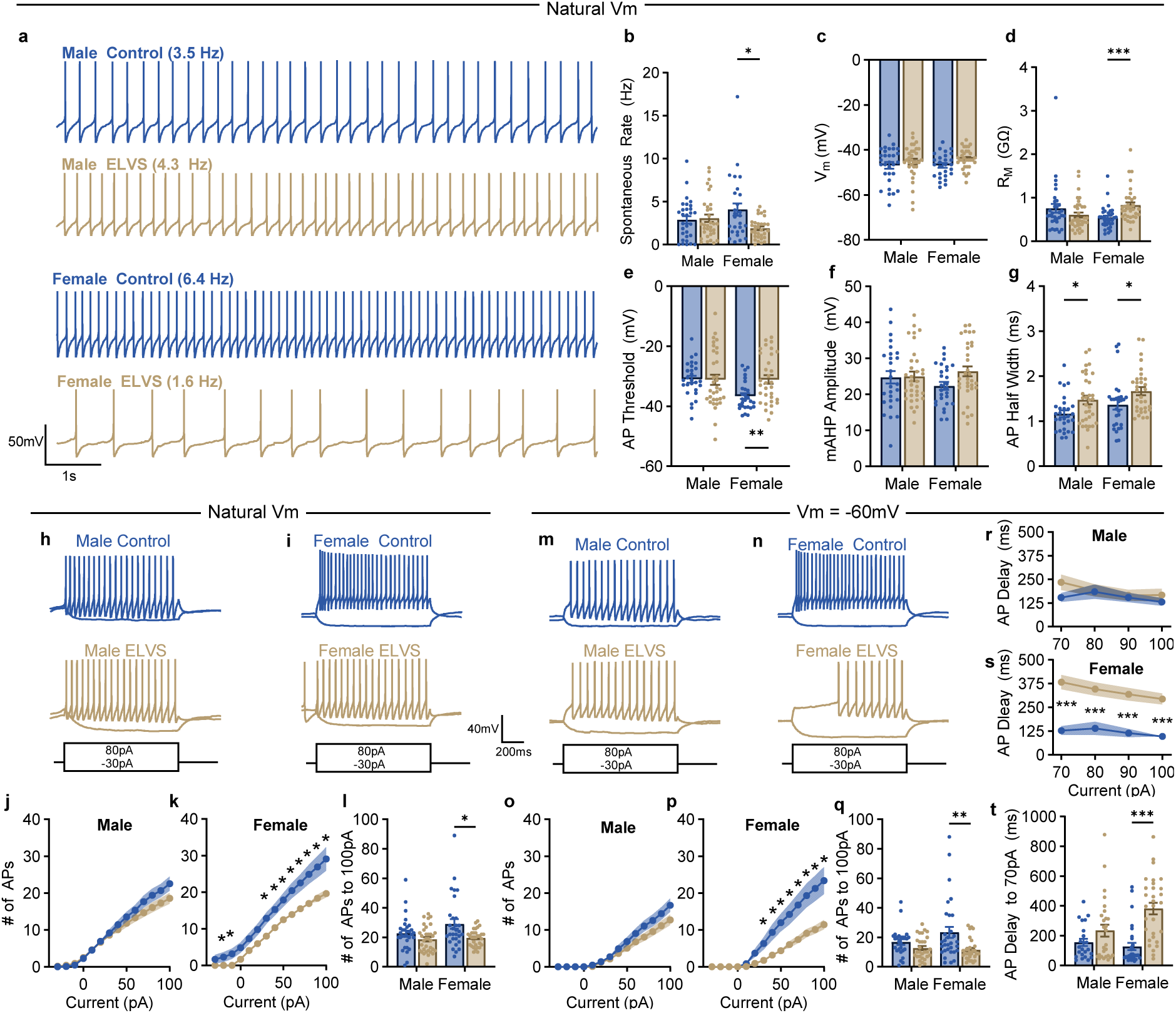
ELVS causes reduced LC neuron excitability and increased AP delay time in female mice. (**a, h, i, m, n**) Representative traces of male and female control and ELVS putative LC neurons when firing spontaneously (**a**), in response to +80/-30 pA current injection (1 s) when cells are at their natural resting membrane potential (**h**, **i**) or at -60 mV (**m**, **n**). Note that the same example recordings for both Vm states are provided from the same neuron, some of which are the same in both resting potential states because the natural Vm was at -60 mV. (**b**-**g**) Action potential spontaneous rate (**b**; males: U = 479.5, *p* = 0.82; females: U = 271.0, p=0.014), resting membrane potential (**c**), membrane resistance (**d**; males: U = 443.0, *p* = 0.14; females: U = 253.5, *p* < 0.0001), action potential (AP) threshold (**e**; males: U = 410.0, *p* = 0.61; females: U = 256.0, *p* = 0.0069), medium after hyperpolarization (mAHP) amplitude (**f**), and AP half-width (**g**; males: U = 314.0, *p* = 0.020; females: U = 295.0, *p* = 0.023) for neurons from each group. (**j**, **k**, **o**, **p**) Number of action potentials elicited by putative LC neurons in response to 1 sec current injection steps from -30 to +100 pA for male (**j**, **o**) and female (**k**, **p**) mice with inter-step membrane potential varying naturally (**j**, **k**; females: U = 293.0 – 370.0, *p* = 0.00030 – 0.031) or maintained near -60 mV (**o**, **p**; females: U = 253.0 – 322.0, *p* = 0.0019 – 0.038). (**l, q**) Number of action potentials elicited by putative LC neurons in response to maximum 100 pA for male and female mice with inter-step membrane potential varying naturally (**l**; males: U = 403.5, *p* = 0.11; females: U = 310.0, *p* = 0.025) or maintained near -60 mV (**q**; males: U = 284.0, *p* = 0.12; females: U = 253.0, *p* = 0.0019). (**r**, **s**) Delay time to first action potential after current injection onset present in putative LC neurons in response to 1 sec current injection steps from +70 to +100 pA for male (**r**) and female (**s**; U = 118.0 – 148.5, *p* < 0.0001) mice with inter-step membrane potential maintained near -60 mV. (**t**) Delay time to first action potential after current injection onset present in putative LC neurons in response to 70 pA current injection (1 sec) for male (U = 258.0, *p* = 0.31) and female (U = 124.0, *p* < 0.0001) mice. All group data are presented as mean ± SEM (bars or shading) for each group, with circles representing data from individual neurons. Two-or three-way ANOVAs were used for group comparisons, followed by a t-test or Mann-Whitney U test for single pairwise comparisons to assess stress effects. n = 26-35 neurons from N=4-5 mice (7-8 weeks old) for each group. For **b**-**g**, **l**, **q**, **r**-**t**, \**p* < 0.05, \*\**p* < 0.01, \*\*\**p* < 0.001 with two-tailed unpaired t-test or Mann-Whitney U test. For clarity in **j**, **k**, **o**, **p**, only \**p* < 0.05 with Mann-Whitney U test for each current injection.

Since the resting potential of LC neurons was quite variable, in the same population of neurons we also recorded evoked action potential firing when constant current was injected to maintain the membrane potential at -60 mV to identify specifically how excitability may change when controlling for V_m_. From an initial resting potential of -60 mV in all neurons, ELVS females continued to show reduced evoked excitability (**Figure 4o-q**), while absent in males (**Figure 4o**) this excitability change appeared to be due to an increase in the delay time to the first action potential following current injection in females (**Figure 4s & t**), at least in part. Although the delay time to the first action potential was variable in all groups, almost all LC neurons in female ELVS mice displayed a notable delay (>40-50 ms) in response to 70 pA current injection, while LC neurons from animals in all other groups displayed both no delay and long delays (**Figure 4t**). This highlights a female-specific LC physiology change that could contribute to the observed sexually dimorphic LC-involved behavioral differences, potentially due to changes in distal release of NE throughout the brain.

To determine the mechanism involved in reducing excitability and increased action potential delay time in ELVS female LC neurons, we first investigated the role of ATP-sensitive K+ channels, which could account for changes in excitability, by removing both cyclic nucleotides (ATP and GTP) from the internal solution to promote maximal K_ATP_ activity and reduce variations in activity that could alter membrane resistance and V_m_ (**Supp.** Figure 7). Although removing ATP altered some stress effects on certain action potential properties, such as the action potential threshold, the medium after hyperpolarization (mAHP) amplitude, and the action potential halfwidth, female ELVS LC neurons remained less excitable both at their resting potential (**Supp.** Figure 4i, **k**, **l**) and at -60 mV (**Supp.** Figure 4n, **p**, **q**). Importantly, the sex-specific ELVS-induced increase in action potential delay time also persisted (**Supp.** Figure 7r-t). It is worth noting the ELVS males showed a significant reduction in spontaneous firing rate when compared to controls when ATP and GTP were excluded, but did not show any significant differences in evoked firing at rest or at -60 mV (**Supp.** Figure 7).

Since action potential delays and suppression of firing rate in response to depolarizing current are due to the activation of transient A-type voltage-gated potassium (K_v_) channel-mediated currents (*I_A_*) known to be present in LC neurons and modifiable [55–57], we pharmacologically assessed the *I_A_*-dependence of this delay with 4-aminopyridine (4-AP, 1 mM). To determine if this difference in A-type potassium channels contributes to the increase in action potential delay time in ELVS females, we recorded current-injection evoked action potentials from a V_m_ of -60 mV before and after bath application of 4-AP (1 mM) to block these transiently activated channels (**Figure 5a**). In nearly all neurons from each group, 4-AP significantly reduced the action potential delay time (**Figure 5b-f**), except for male ELVS mice where the delay time was already short. In the presence of 4-AP, the action potential delay times were comparable in all groups (**Figure 5f**). These data indicate that the 4-AP-sensitive increased action potential delay time in ELVS female LC neurons is due to some change in expression or modulation of the K_v_ channel or accessory protein mediating *I_A_* that occurs due to ELVS exposure.

**Figure 5.**
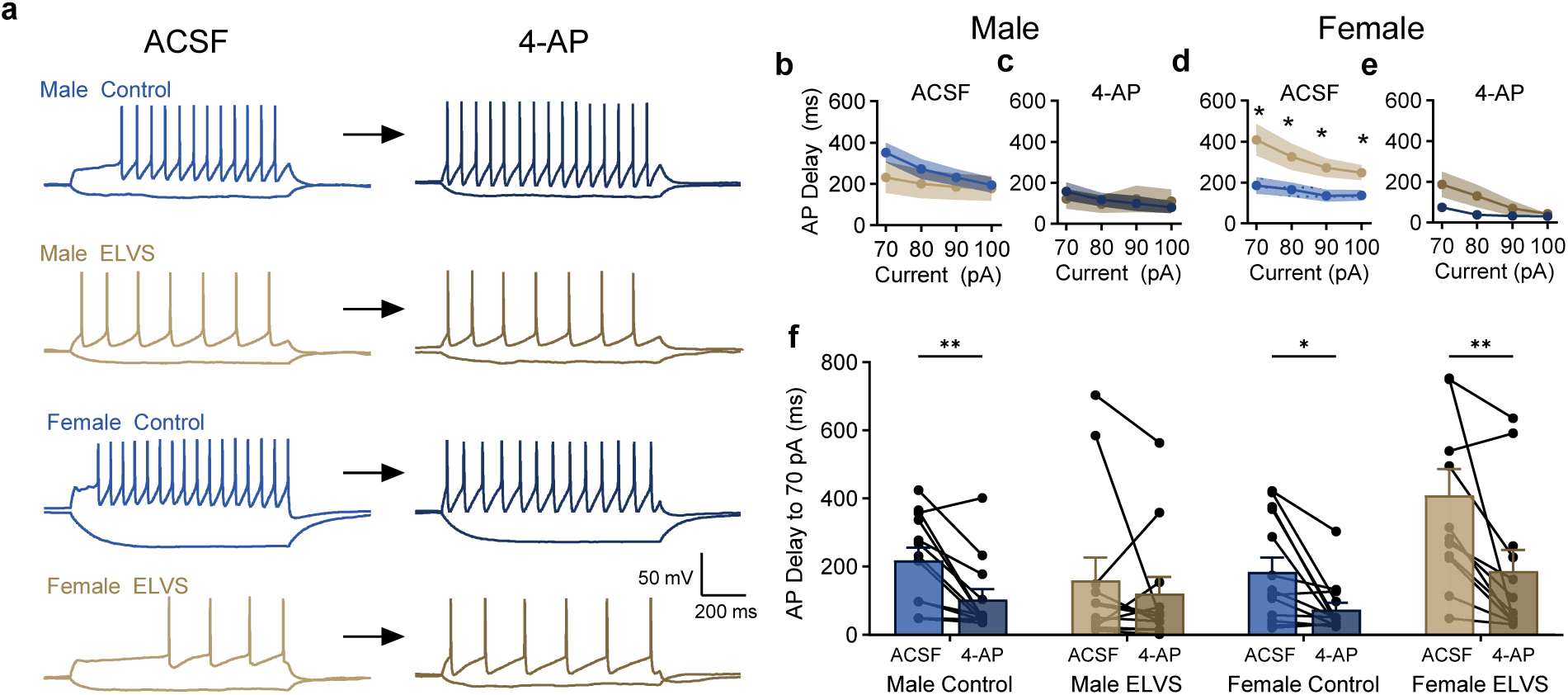
Blockade of K_A_ (A-type) potassium channels reduces action potential delay time and normalizes ELVS-induced action potential delay time prolongation. (**a**) Representative recordings from LC neurons in male (top) and female (bottom) control and ELVS mice before (left) and after (right) 5 min bath application of 20 mM 4-aminopyridine (4-AP). **b**-**e**) Delay time to first action potential after current injection onset present in putative LC neurons in response to 1 sec current injection steps from +70 to +100 pA before (**b**, **d**) and after 4-AP application (**c**, **e**) for male (**b**, **c**) and female (**d**,**e**; control: U = 40.0 – 50.0, *p* = 0.023 – 0.047; ELVS: U = 64.0 - 80.0, *p* = 0.47 – 0.84) control (blue) and ELVS (gold) mice with inter-step membrane potential maintained near -60 mV. (**f**) Delay time to first action potential after current injection onset present in putative LC neurons in response to 70 pA current injection (1 sec) for male (control: *W* = 80, *p* = 0.0024; ELVS: *W* = 24, *p* = 0.38) and female (control: *W* = 79, *p* = 0.010; ELVS: *W* = 74, *p* = 0.0015) mice. All group data are presented as mean ± SEM (bars or shading) for each group, with connected circles representing paired data from individual neurons. Three-way ANOVAs and repeated measures ANOVAs were used for group comparisons, followed by a Mann-Whitney U test for single pairwise comparisons to assess stress effects or Wilcoxon Rank-Sum for paired 4-AP effects. n = 11-14 neurons from N=3 mice (7-8 weeks old) for each group. \**p* < 0.05, \*\**p* < 0.01 for each indicated pairwise comparison.

### Restoration of extracellular NE levels by reboxetine ameliorated ELVS-induced changes in behavior

Electrophysiological recordings of LC neurons indicate a reduced excitability and increased delay to respond to current injection, both of which would lead to reduced distal NE release. Therefore, to determine the impact of these physiological changes on behavior, we assessed whether increasing norepinephrine levels at noradrenergic synapses using reboxetine, a selective norepinephrine reuptake inhibitor, would restore behaviors found to be altered in separate cohorts of control and ELVS mice (**Figure 6**). We found that 2mg/kg oral reboxetine administration restored female-specific effects of ELVS on sucrose preference behavior (**Figure 6a** & **b**). Oral reboxetine also significantly reduced exploratory behavior in ELVS mice (**Figure 6c** & **d**), but did not alter y-maze performance (**Figure 6e** & **f**). Overall, this indicates that manipulating the LC-NE system in a manner that is targeted based on the sex- and stress-dependent changes in physiology may be beneficial in restoring specific affective behavioral changes.

**Figure 6.**
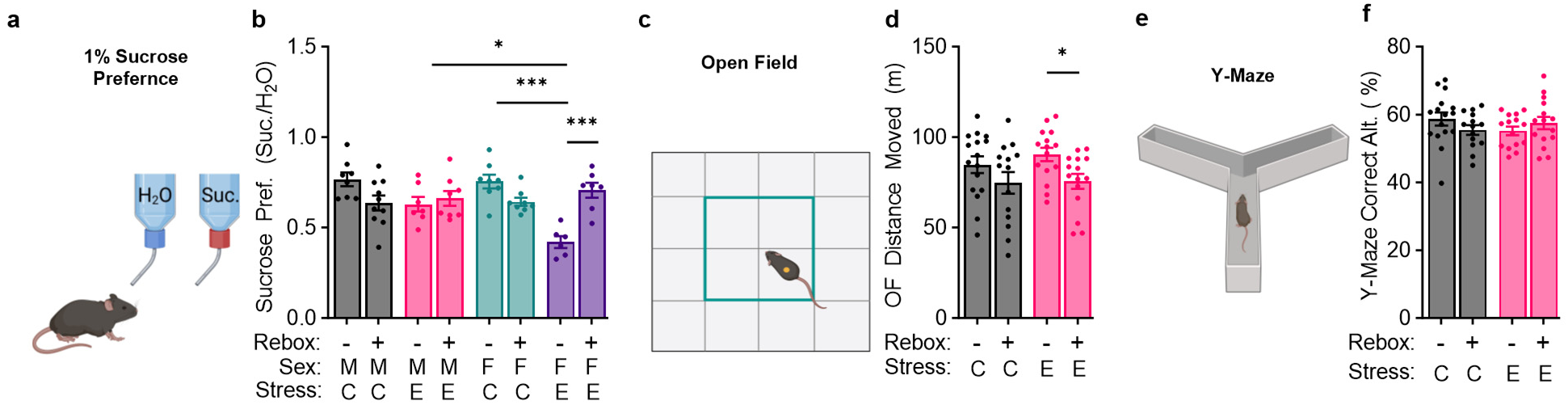
Norepinephrine transporter blockade ameliorates ELVS-induced reduced sucrose preference and increased exploratory activity in female mice. Performance of control and ELVS mice after oral consumption of vehicle (apple juice) or the norepinephrine transporter inhibitor reboxetine (2 mg/kg) in the sucrose preference (**a**, **b**; ELVS female vehicle vs. reboxetine: *p* = 0.0001; Control vs. ELVS female: *p* < 0.0001; ELVS male vs. female: *p* = 0.010), open field (**c**, **d**; control vehicle vs. reboxetine: U = 79.0, *p* = 0.18; ELVS vehicle vs. reboxetine: U = 52.0, *p* = 0.011), or y-maze (**e**, **f**) assays. All group data are presented as mean ± SEM (bars) for each group, with circles representing data from individual mice. Three- or two-way ANOVAs were used for group comparisons, followed by a Bonferroni posthoc or Mann-Whitney U test for pairwise comparisons to assess effects of oral reboxetine treatment on group performance. N = 6-10 mice (7-9 weeks old) for each group and sex. \**p* < 0.05, \*\*\**p* < 0.001 for pairwise comparisons.

## Discussion

### Summary of Findings

In our two-phase animal model developed to mimic ACE-relevant timing of and systematic variation in exposure to threat and deprivation in early life in mice, we identified altered exploratory locomotion in a familiar environment, short-term memory in early adulthood, and evidence of reduced ability to maintain attention, regardless of sex (**Figures 1 & 2**). We also found male-specific increases in impulsivity and evidence of female-specific reduced impulsivity, increased exploratory behavior, and an anhedonia-like state (**Figures 1 & 2**). Notably, changes in basal anxiety-like behaviors were absent due to ELVS exposure in the B6J mice used here, in contrast to Balb/C mice used in our previous work [58], where a prominent increase in anxiety-like behavior was observed in female mice. Motivated by the role of norepinephrine release from the LC in shaping these behaviors, our evaluation of LC neurons in these mice using electrophysiological recordings identified a reduction in LC neuron excitability that was due in part to a prolongation of the onset action potential delay (**Figure 4**), which was mediated by a 4-AP-sensitive A-type Kv channel (**Figure 5**). Unlike male Balb/C mice exposed to a similar ELVS protocol [58], basal properties of LC neurons from B6J male mice exposed to ELVS were remarkably similar to those from control mice. In addition to blunted CRF sensitivity specifically in female mice exposed to ELVS, in LC neurons from naïve mice, CRF-induced excitability changes were due to involvement of sex-specific signaling pathways (**Figure 3**), which may partly explain why exposure to stressful stimuli during adolescence leads to sex-specific changes in basal LC neuron excitability. Finally, pharmacologically compensating for this loss of LC neuron excitability/NE release by blocking NE reuptake with reboxetine [59] restored a subset of ELVS-induced behavioral changes (**Figure 6**), a drug that has been used clinically to treat depression, panic disorder, and ADHD [60–62] with variable success. Together, these data highlight the dynamics and importance of understanding the complex interplay between sex, age (critical periods), genotypic variation, stress exposure, and neuromodulator circuit changes.

### The Impact of Stress and Sex on LC Function Related to Affect and Cognition

The LC-NE system is also well-documented to be stress-sensitive, as it is activated by an array of environmental stressors, including social and predator stress [19, 21, 63–69]. Largely limited to stress exposure in males, acute stress-induced changes in LC_NE_ activity is proposed to occur primarily through the activation of CRFR_1_ by CRF released from several brain areas, including the central amygdala and paraventricular nucleus of the hypothalamus [19, 64, 70–73]. At least in male rats, stress and stress-related CRF release shift LC activity towards a high tonic firing mode (3-8 Hz) while simultaneously decreasing phasic firing events [72, 74, 75]. These stress-induced increases in CRF and modulation of LC-NE neuron activity are paralleled by CRF-induced changes in multiple processes and behaviors known to be heavily shaped by LC-NE neuron activity, including attention [76], cognitive flexibility [19], anxiety [10, 15], and memory [77]. The data presented here shed light on how the LC-NE system changes with multiple early life stress exposures, and likely related increase in CRF release, depending on sex.

The nature of the LC physiology changes of female mice due to ELVS exposure has the potential to affect both tonic and phasic firing, each of which is proposed to be related to specific behavioral events [16, 17, 28, 78]. First, the reduced LC spontaneous and persistent evoked firing rate observed only in ELVS-exposed females will affect tonic firing. Second, by reducing the likelihood of generating an action potential in response to short depolarizing events (i.e. the beginning of current injection steps), the prolongation of the action potential delay time will act as a filter of fast synaptic inputs to reduce the likelihood of phasic firing and subsequent distal NE release. By disrupting both types of LC neuron firing modes, these changes observed in females could result in disrupted NE signaling in multiple brain regions involved in attention, arousal, memory, and/or locomotion (i.e. prefrontal cortex, hippocampus, ventral tegmental area, and lateral cerebellar nucleus).

Although LC-NE neurons were previously thought to be a homogenous population, recent data suggest that they are much more heterogeneous in their molecular [79] or functional properties that relate to specific LC-NE projection targets [80, 81]. Given the range of delay times, resting potentials, and spontaneous firing rates observed in this population of LC neurons, it appears that ELVS exposure leads to a shift in the distribution of these key properties across the population, likely due to sex-specific variation in CRF-activated signaling. However, it is not clear how a change in these functional properties may relate to changes in specific molecularly-, anatomically-(target brain area), or functionally-defined LC neuron sub-populations. Whether acute or persistent changes in the physiological properties of LC-NE neurons and/or their synaptic properties, like those observed here, may impact NE release throughout the central nervous system differentially based on specific circuits or related behavior is unclear. Since there is a smaller distribution of firing rates yet lager distribution of action potential delay times across LC neurons in ELVS females when compared to controls, this property-dependent shift in the functional diversity of LC neurons could impact how firing modes required for optimal projecting brain region states. For instance, distinct LC ensembles projecting to the cortex may modulate particular cortical states with members of each ensemble having definitive firing rates [82]. Further effort to determine exactly how these changes in LC-NE system excitability alter widespread noradrenergic signaling *in vivo* during specific behaviors would provide support for the importance of this physiological change and suggest the LC-NE system to be a female-specific target of interest for stress-related disorders, particularly with onset after adolescence.

The specific sexually dimorphic molecular mechanisms underpinning LC physiology changes after exposure to the ELVS are still unclear. However, our results in naïve mice (**Figure 3**) establish that there are basal sex differences in how CRF exposure affects the firing of LC neurons, at least in the signaling pathways involved in CRF-induced increases in firing rate, indicating preferential G_q_ signaling in males and G_s_ in females. These data are in line with other reports of female CRFR1 in the LC being predominantly G_s-_coupled and males having increased β-arrestin-related internalization of CRFR1 after acute stress [32]. Bridging the gap in future work to understand how two-phase early life stress and acute stress affect CRF-dependent changes in LC physiology and related signaling based on sex will be important to understand fundamental sex-specific processes that shape responses to stress on both timescales. Based on these data, we predict that the predominant G_s_-dependent CRF actions in females, leading to PKA activation, will cause unique subsequent and persistent changes in LC physiology (excitability, delay) in females that will be absent in males. In parallel, the G_q_ coupling predominance could be protective in males due to its transient Ca^2+^-dependent action versus a prolonged upregulation of a signaling cascade. Given recent reports of persistent LC physiology changes occurring only when stress exposure happens at adolescence and not adulthood [83], determining the degree to which this applies to our dual-phase ELVS paradigm and the duration of any lasting behavioral and physiological changes will also be important to understand dynamics that shape the broader impacts of stress. Finally, at a molecular level, it is unclear how exactly *I_A_* is modulated or affected by stress, but clarifying the channels and auxiliary subunit changes over longer time scales may reveal novel mechanisms of broader and LC-specific responses to critically-timed stress to shape novel prevention and treatment strategies to improve specific behavioral outcomes.

### Clinical Impact of Findings

The treatment of stress- or specifically ACE-related disorders should consider sex and the timing of stress exposure as biological variables influencing patient outcomes. These behavioral results suggest females may be more sensitive to depression-like behaviors and ADHD-relevant deficits as a result of early life stress during comparable time periods used in this study. Based on these data, individuals exposed to ACEs as adolescents may be particularly sensitive to developing negative neuropsychiatric outcomes later in life, based on sex differences in LC neuron physiology and central stress (CRF) sensitivity at this time. Regardless, the specific role of sex hormones in development and neurophysiological changes in response to stress is unclear. Estrogen receptor α (ERα) and β (ERβ) are differentially expressed in the amygdala, prefrontal cortex, and hippocampus responsible for stress, anxiety, and cognitive behaviors, respectively [84]. ERβ has been associated with non-reproductive behaviors, with its activation improving memory and reducing anxious behaviors in females [85, 86]. In contrast, testosterone signaling has been suggested to be a factor supporting resilience to stress-induced changes, which is in line with the behavioral data presented here [87, 88].

This study may implicate the adrenergic system, particularly the LC in the brain, as a region to explore biomarkers of dysfunction in female patients with a history of stress in early life. According to the DSM-V, cognitive issues are a core symptom of depression that include aspects of memory and attention being disrupted [89, 90]. These symptoms are in line with the female-specific behavioral impacts of our dual-phase stress model, pointing to increasing noradrenergic tone as a therapeutic avenue for treatment. There are many drugs that target this brain area, particularly second-line therapies for attention deficit hyperactivity disorder (ADHD), such as reboxetine [61] used in this study, which may prove to be effective in managing cognitive impairments in this patient group with depression and/or PTSD. These data provide a foundation for further research into neuro-stress regulation differences in males and females, relevant to many stress-related disorders, from PTSD to substance abuse disorder, that have sex differences in their prevalence ratios [46]. Perhaps a more robust individualized understanding of how patients’ genetic and environmental risk factors interact with the type of timing of major stressful life events may increase the likelihood of therapeutic success when these variables are used to optimize drug choice.

## Acknowledgements

This work was supported by institutional start-up funds and a Research Seed Grant from SIU-School of Medicine provided to BDR.

