## Supplemental Information for "Early life stress induces sex-specific changes in behavior and parallel locus coeruleus neuron excitability"

### Supplementary Information

### Supplementary Figure 1 – NOR/Rotarod

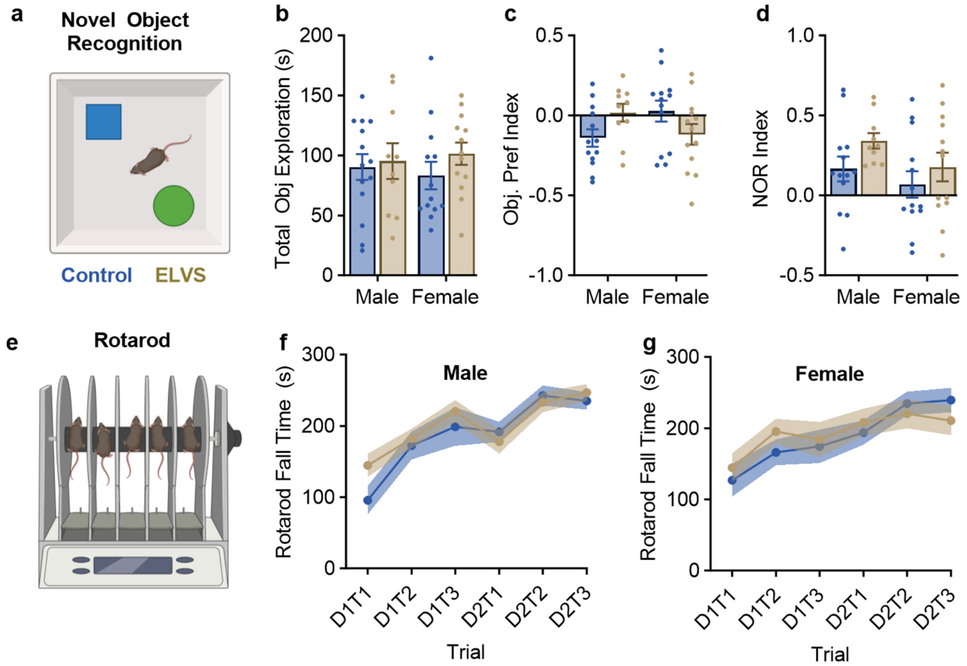


**Supplementary Figure 1. ELVS exposure does not affect long-term (24hr) memory or motor function.** (**a-d**) Novel Object Recognition (NOR) assay (**a**) total cumulative object exploration time (**b**), initial object preference index in the second phase (**c**), and NOR index in the third phase (**d**). (**e-g**) Two-day rotarod assay latency to fall times for each of three trials from both days in male (**f**) and female (**g**) control (blue) and ELVS (tan) mice. All results are presented as mean ± SEM for each group with circles representing data from individual mice. N = 9-14 mice (6 weeks) per group.

### Supplementary Figure 2 – Full 5CSRTT

**a**

- Test 1: Long ITI
- Test 2: Short ITI
- Test 3: Reduced stimulus duration
- Test 4: Sound distractor (1st time)
- Test 5: Light intensity changes (random)
- Test 6: Sound distractor (2nd time)
- Test 7: Sound distractor + light intensity changes
- Test 8: Sound + light + long ITI
- Test 9: Sound + light + short ITI
- Test 10: Sound + light + reduced stimulus duration

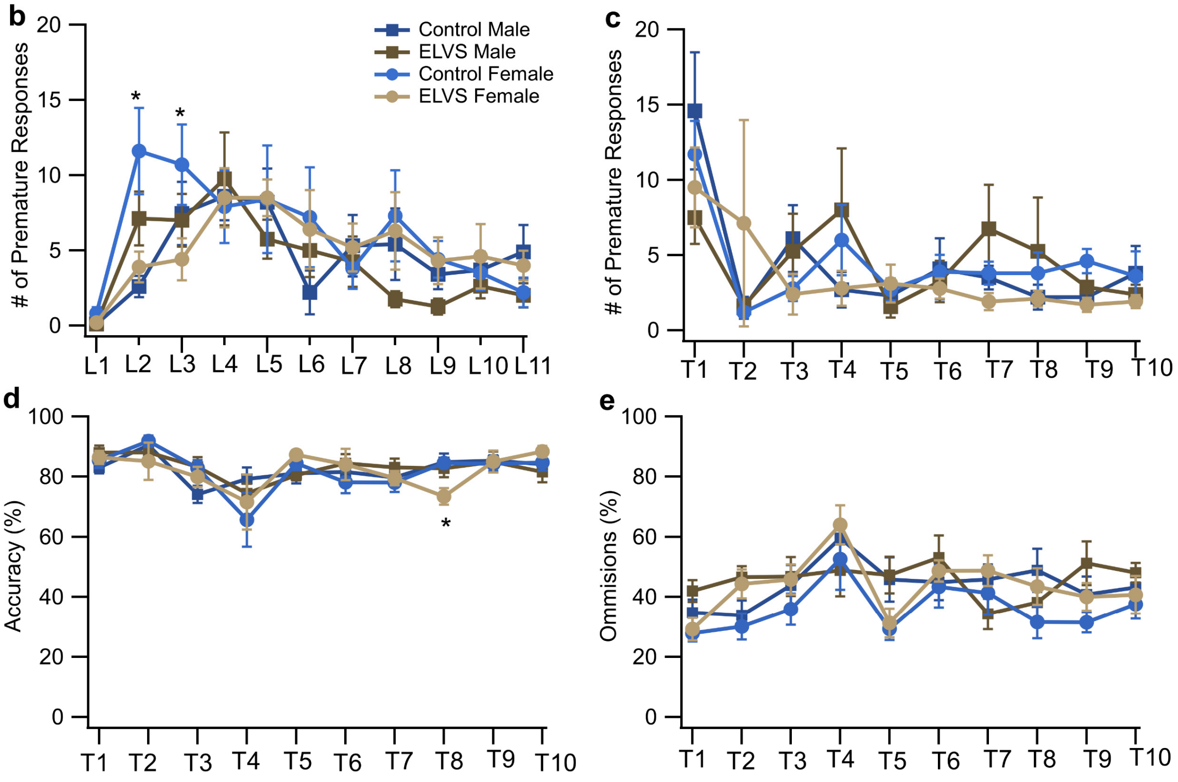


**Supplementary Figure 2. 5SCRTT Premature Responses during training and accuracy and omissions during testing.** (**a**) Listing of session variable changes for each of the ten test sessions. (**b**) Mean ± SEM number of premature trials of each group for the first time performing each training level. (**c-e**) Mean ± SEM number of premature responses (**c**), accuracy % (**d**), and omissions % (**e**) for each group during each test. N=7-10 mice (6-8 months old). * p<0.05 with unpaired two-tailed t-test for each sex for each session.

**Supplementary Figure 3 – LCA & Holding Time**

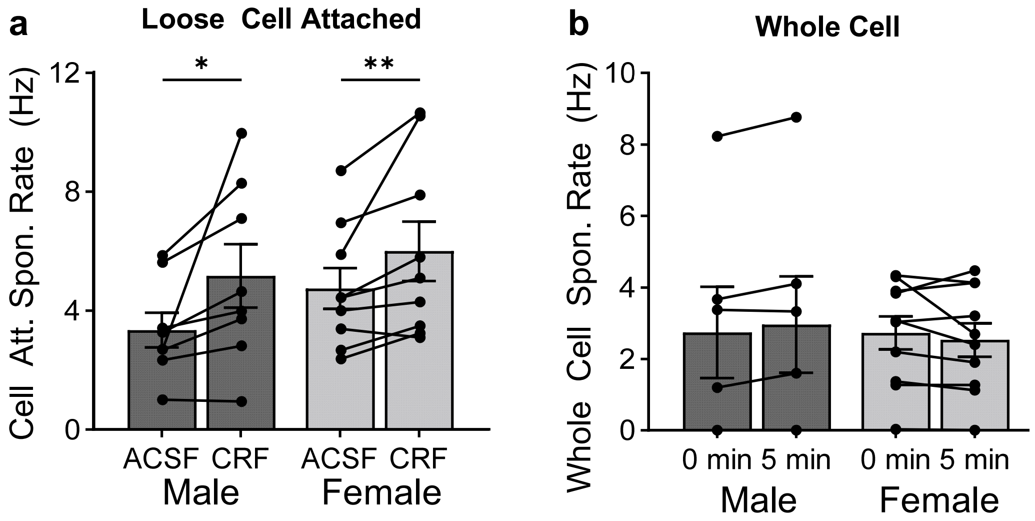


**Supplementary Figure 3. Whole cell recording configuration and recording duration do not impact CRF-induced LC neuron excitability changes.** (**a**) Spontaneous firing rate of putative LC neurons from 5–6-week-old male (W = -34.00, $p = 0.016$) and female (W = -43.00, $p = 0.0078$) mice in a loose cell attached configuration before and 5 min after bath applied 500 nM. (**b**) Spontaneous firing rate (Hz) of putative LC neurons from 5–6-week-old male and female mice recorded in the whole cell configuration at time = 0 min, beginning after a standard 5 min period after obtaining the whole cell configuration and after 5 min of recording. All group data are presented as mean ± SEM (bars or shading) for each group, with connected circles representing paired recording data from individual neurons. Two-way RMANOVAs were used for group comparisons, followed by a Wilcoxon Rank-Sum (paired) test (*$p < 0.05$, **$p < 0.01$) for single pairwise comparisons to assess CRF and time effects. n = 6-10 neurons from N=2-3 mice (5-6 weeks old) for each group.

### Supplementary Figure 4 – CRF & Evoked Firing

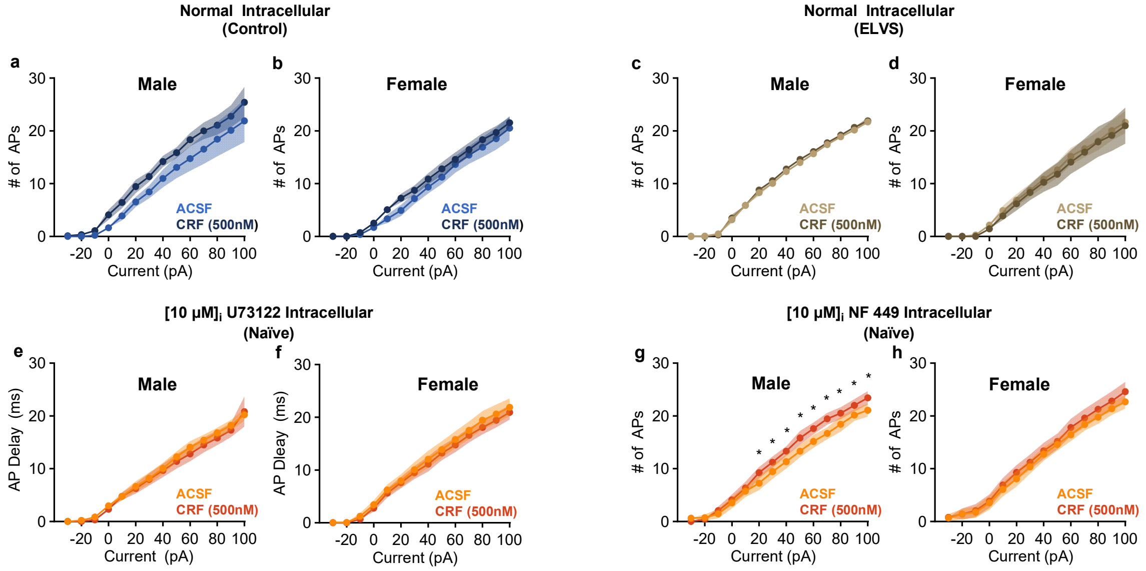


**Supplementary Figure 4. CRF does not significantly impact evoked firing.** (**a-d**) Number of action potentials elicited in response to 1 sec current injection steps (-30 to +100 pA) before and 5 min after bath application of CRF (500nM) by putative LC neurons from male (**a**, **c**) and female (**b**, **d**) control (**a**, **b**) and ELVS-exposed (**c**, **d**) mice. (**e-h**) Number of action potentials elicited in response to 1 sec current injection steps (-30 to +100 pA) before and 5 min after bath application of CRF (500nM) by putative LC neurons from naïve male (**e**, **g**) and female (**f**, **h**) naïve mice with either the phospholipase C inhibitor, U73122 (10μM; **e**, **f**; male: W = -30.00 – -66.00, $p$ = 0.0010 – 0.039), or the G$\alpha_s$ inhibitor, NF 449 (10μM; **g**, **h**), included in the intracellular recording solution. All group data are presented as mean ± SEM (shading) for each group. Two or three-way RMANOVAs were used for group comparisons, followed by a Wilcoxon Rank-Sum (paired) test (*$p$ < 0.05) for single pairwise comparisons to assess CRF effects within each group. n = 9-13 neurons from N = 3 mice (5-7 weeks old) for each group.

**Supplementary Figure 5 – ELVS Synaptic Ephys**

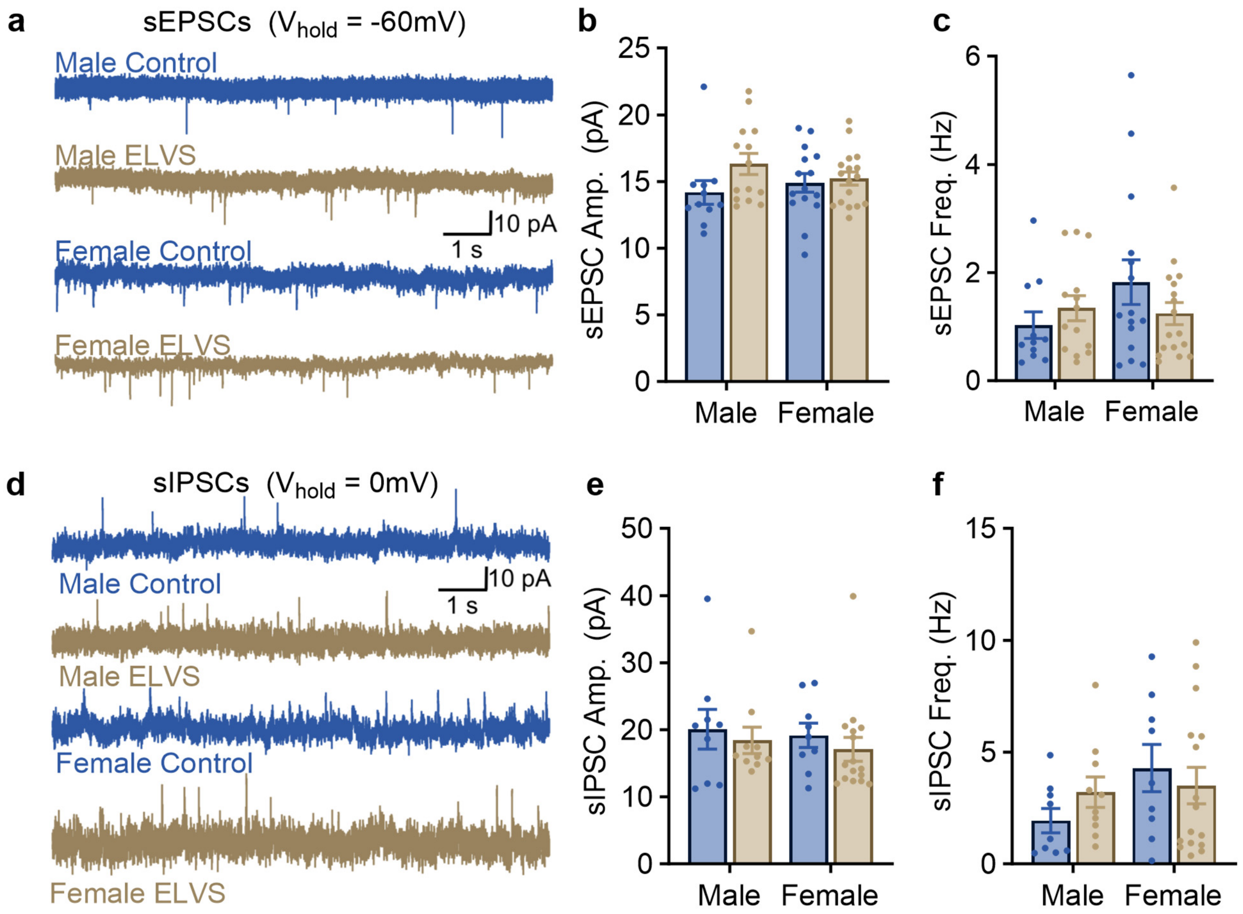


**Supplementary Figure 5. ELVS does not impact LC synaptic physiology.** (**a**, **d**) Representative voltage clamp recordings of male and female control and ELVS putative LC neuron spontaneous excitatory postsynaptic currents (sEPSCs; **a**; ($V_{hold}$ = -60 mV; $E_{Cl}$ = -85 mV) or spontaneous inhibitory postsynaptic currents (sEPSCs; **d**; $V_{hold}$ = 0 mV; $E_{Cl}$ = -85 mV). Group amplitude (**b**, **e**) and frequency (**c**, **f**) of sEPSCS (**b**, **c**) and sIPSCS (**e**, **f**). All group data are presented as mean ± SEM (bars) for each group, with circles representing data from individual neurons. Two-way ANOVAs were used for group comparisons. n = 9-17 neurons from N = 3-4 mice (7-9 weeks old) for each group.

**Supplementary Figure 6 – No Ephys Change Early**

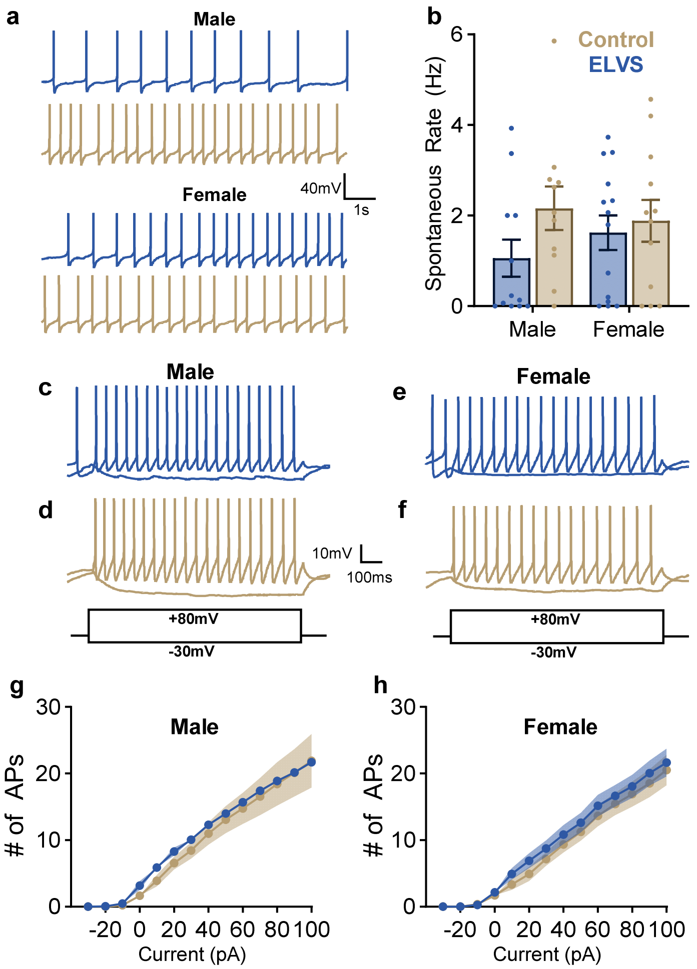


**Supplementary Figure 6. No LC neuron excitability changes immediately after ELVS.** (**a**, **c**, **d**, **e**, **f**) Representative recordings of the membrane potential from putative LC neurons obtained within one week after ending ELVS or control handling with neurons firing action potentials spontaneously (**a**) and from male (**c**, **d**) and female (**e**, **f**) control (**c**, **e**) and ELVS (**d**, **f**) mice in response to +80/-30 pA current injection (1 s) when cells are at their natural resting membrane potential. (**b**, **g**, **h**) Action potential spontaneous rate (**b**) and number of action potentials elicited by putative LC neurons in response to 1 sec current injection steps for male (**g**) and female (**h**) mice with inter-step membrane potential varying naturally. All group data are presented as mean ± SEM (bars or shading) for each group, with circles representing data from individual neurons. Two- or three-way ANOVAs were used for group comparisons. n = 9-13 neurons from N = 3 mice

(6 weeks old) for each group. ANOVAs did not detect significant effects of stress exposure of sex in any cases.

### Supplementary Figure 7 - No ATP/GTP

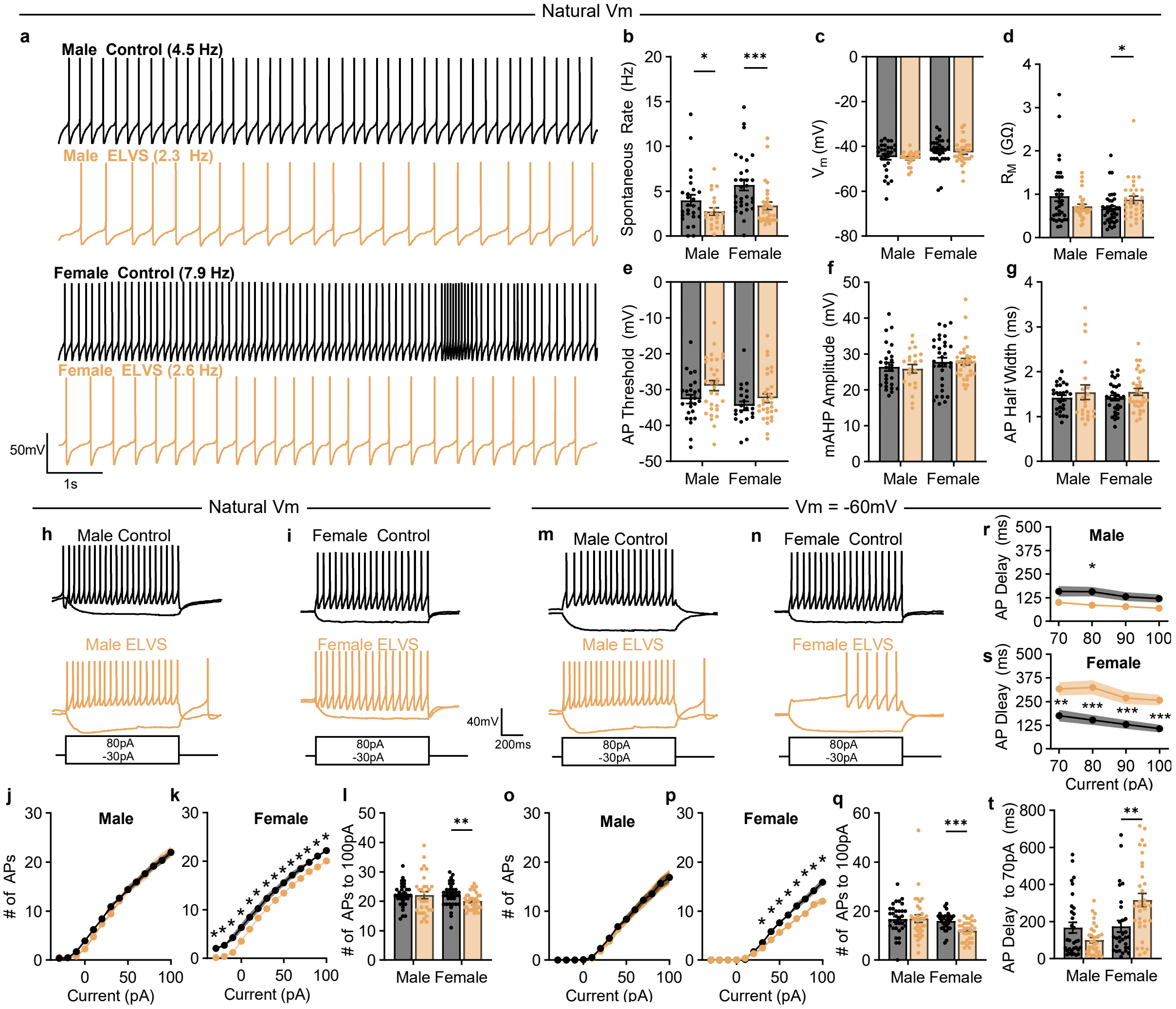


**Supplementary Figure 7. ELVS-induced changes in LC excitability persist despite limiting $K_{ATP}$ activity by removing intracellular ATP/GTP.** All recordings completed with ATP and GTP omitted from the whole cell internal solution. (**a, h, i, m, n**) Representative traces of male and female control and ELVS putative LC neurons when firing spontaneously (**a**) in response to +80/-30 pA current injection (1 s) when cells are at their natural resting membrane potential (**h**, **i**) or at -60 mV (**m**, **n**). Note that the same example recordings for both Vm states are provided from the same neuron, some of which are the same in both resting potential states because the natural Vm was at -60 mV. (**b-g**) Action potential spontaneous rate (**b**; males: U = 174.0, *p* = 0.039;

females: U = 231.5, p=0.0007), resting membrane potential (**c**), membrane resistance (**d**; males: U = 460.5, *p* = 0.51; females: U = 401.5, *p* = 0.021), action potential (AP) threshold (**e**; males: U = 292.0, *p* = 0.077; females: U = 272.0, *p* = 0.33), medium after hyperpolarization (mAHP) amplitude (**f**), and AP half-width (**g**) for neurons from each group. (**j**, **k**, **o**, **p**) Number of action potentials elicited by putative LC neurons in response to 1 sec current injection steps from -30 to +100 pA for male (**j**, **o**) and female (**k**, **p**) mice with inter-step membrane potential varying naturally (**j**, **k**; females: U = 341.5 – 408.5, *p* = 0.0021 – 0.021) or maintained near -60 mV (**o**, **p;** females: U = 225.0 – 354.5, *p* = 0.00012 – 0.049). (**l, q**) Number of action potentials elicited by putative LC neurons in response to maximum 100 pA for male and female mice with inter-step membrane potential varying naturally (**l**; males: U = 460.0, *p* = 0.63; females: U = 367.0, *p* = 0.0061) or maintained near -60 mV (**q**; males: U = 475.5, *p* = 0.50; females: U = 225.0, *p* = 0.0001). (**r**, **s**) Delay time to first action potential after current injection onset present in putative LC neurons in response to 1 sec current injection steps from +70 to +100 pA for male (**r**; U = 337.0, *p* = 0.019) and female (**s**; U = 211.0 – 281.0, *p* = 0.0001 – 0.029) mice with inter-step membrane potential maintained near -60 mV. (**t**) Delay time to first action potential after current injection onset present in putative LC neurons in response to 70 pA current injection (1 sec) for male (U = 422.5, *p* = 0.23) and female (U = 281.0, *p* = 0.0029) mice. All group data are presented as mean ± SEM (bars or shading) for each group, with circles representing data from individual neurons. Two- or three-way ANOVAs were used for group comparisons, followed by a t-test or Mann-Whitney U test for single pairwise comparisons to assess stress effects. n = 20-37 neurons from N = 4-5 mice (7-8 weeks old) for each group. For **b-g**, **l**, **q**, **r-t**, *$p < 0.05$, **$p < 0.01$, ***$p < 0.001$, and for clarity in **j**, **k**, **o**, **p**, only *$p < 0.05$ with Mann-Whitney U test for each pairwise comparison.
