## Supplemental Table 1 for "Early life stress induces sex-specific changes in behavior and parallel locus coeruleus neuron excitability"

| Data | Group Comparison | n (cells) | N (animals) | Spearman's Heteroscedasticity | Shapiro-Wilk Normality p-value | Term | Statistic | p-value | Pairwise test |
|---|---|---|---|---|---|---|---|---|---|
| Figure 1b | 3-Way RMANOVA | | 16, 16, 18, 17 | **0.0012** | **0.0004** | Time | F (5, 378) = 53.28 | **<0.0001** | Mann-Whitney |
| | | | | | | Sex | F (1, 378) = 1.398 | 0.2377 | |
| | | | | | | Stress | F (1, 378) = 29.30 | **<0.0001** | |
| | | | | | | Time x Sex | F (5, 378) = 1.172 | 0.3221 | |
| | | | | | | Time x Stress | F (5, 378) = 0.7151 | 0.6124 | |
| | | | | | | Sex x Stress | F (1, 378) = 14.40 | **0.0002** | |
| | | | | | | Time x Sex x Stress | F (5, 378) = 0.08822 | 0.9941 | |
| Figure 1c | 2-Way ANOVA | | 16, 16, 18, 17 | 0.3947 | 0.3286 | Sex x Stress | F (1, 63) = 3.771 | **0.0566** | Unpaired t-test |
| | | | | | | Sex | F (1, 63) = 0.4632 | 0.4986 | |
| | | | | | | Stress | F (1, 63) = 9.355 | **0.0033** | |
| Figure 1d | 2-Way ANOVA | | 13, 10 13, 13 | 0.3726 | 0.6074 | Sex x Stress | F (1, 45) = 1.110 | 0.2978 | Unpaired t-test |
| | | | | | | Sex | F (1, 45) = 0.1536 | 0.6970 | |
| | | | | | | Stress | F (1, 45) = 8.956 | **0.0045** | |
| Figure 1e | 2-Way ANOVA | | 16, 16, 18, 17 | 0.0827 | 0.5966 | Sex x Stress | F (1, 63) = 4.854 | **0.0313** | Unpaired t-test |
| | | | | | | Sex | F (1, 63) = 0.6763 | 0.4140 | |
| | | | | | | Stress | F (1, 63) = 10.51 | **0.0019** | |
| Figure 1f | 2-Way ANOVA | | 16, 16, 18, 17 | 0.4497 | **0.0171** | Sex x Stress | F (1, 63) = 0.9145 | 0.3426 | None |
| | | | | | | Sex | F (1, 63) = 1.324 | 0.2542 | |
| | | | | | | Stress | F (1, 63) = 0.9814 | 0.3256 | |
| Figure 1h | 2-Way ANOVA | | 15, 16, 18, 16 | **0.0192** | 0.2724 | Sex x Stress | F (1, 61) = 3.143 | 0.0813 | None |
| | | | | | | Sex | F (1, 61) = 1.946 | 0.1681 | |
| | | | | | | Stress | F (1, 61) = 2.903 | 0.0935 | |
| Figure 1j | 2-Way ANOVA | | 16, 14, 18, 16 | 0.4328 | 0.6450 | Sex x Stress | F (1, 61) = 0.7598 | 0.3868 | Unpaired t-test |
| | | | | | | Sex | F (1, 61) = 0.06325 | 0.8023 | |
| | | | | | | Stress | F (1, 61) = 8.149 | **0.0059** | |
| Figure 1k | 2-Way ANOVA | | 16, 14, 18, 16 | 0.3030 | 0.8005 | Sex x Stress | F (1, 61) = 2.049 | 0.1574 | Unpaired t-test |
| | | | | | | Sex | F (1, 61) = 0.01929 | 0.8900 | |
| | | | | | | Stress | F (1, 61) = 7.978 | **0.0064** | |
| Figure 1m | 2-Way ANOVA | | 8, 8, 8, 8 | 0.2763 | 0.3659 | Sex x Stress | F (1, 28) = 11.76 | **0.0019** | Unpaired t-test |
| | | | | | | Sex | F (1, 28) = 21.88 | **0.0001** | |
| | | | | | | Stress | F (1, 28) = 33.95 | **0.0000** | |
| Figure 2a | 2-Way ANOVA | | 10, 8, 10, 10 | **0.0030** | 0.2067 | Sex x Stress | F (1, 34) = 0.7207 | 0.4019 | None |
| | | | | | | Sex | F (1, 34) = 5.294 | **0.0277** | |
| | | | | | | Stress | F (1, 34) = 1.050 | 0.3128 | |
| Figure 2b | 2-Way ANOVA | | 10, 8, 10, 10 | 0.2392 | 0.2988 | Sex x Stress | F (1, 34) = 0.3173 | 0.5769 | Unpaired t-test |
| | | | | | | Sex | F (1, 34) = 8.011 | **0.0078** | |
| | | | | | | Stress | F (1, 34) = 4.285 | **0.0461** | |
| Figure 2c | 2-Way ANOVA | | 10, 8, 10, 10 | **0.0001** | 0.6839 | Sex x Stress | F (1, 34) = 12.58 | **0.0012** | Mann-Whitney |
| | | | | | | Sex | F (1, 34) = 2.161 | 0.1507 | |
| | | | | | | Stress | F (1, 34) = 1.049 | 0.3130 | |
| Figure 2d | 2-Way ANOVA | | 10, 8, 10, 10 | 0.4321 | **0.0066** | Sex x Stress | F (1, 34) = 4.054 | 0.0592 | None |
| | | | | | | Sex | F (1, 34) = 0.8250 | 0.0520 | |
| | | | | | | Stress | F (1, 34) = 1.022 | 0.3701 | |
| Figure 2e | 2-Way ANOVA | | 10, 8, 10, 10 | **0.0071** | **<0.001** | Sex x Stress | F (1, 34) = 0.3114 | 0.5805 | None |
| | | | | | | Sex | F (1, 34) = 0.04442 | 0.8343 | |
| | | | | | | Stress | F (1, 34) = 1.414 | 0.2427 | |
| Figure 2f | 2-Way ANOVA | | 10, 8, 10, 10 | 0.3325 | **0.0053** | Sex x Stress | F (1, 34) = 3.090 | **0.0878** | Mann-Whitney |
| | | | | | | Sex | F (1, 34) = 3.453 | 0.0718 | |
| | | | | | | Stress | F (1, 34) = 6.460 | **0.0158** | |
| Figure 2g | 2-Way ANOVA | | 10, 8, 10, 10 | 0.2645 | 0.3023 | Sex x Stress | F (1, 34) = 0.3426 | 0.5622 | None |
| | | | | | | Sex | F (1, 34) = 0.5671 | 0.4566 | |
| | | | | | | Stress | F (1, 34) = 1.181 | 0.2848 | |
| Figure 2h | 2-Way ANOVA | | 10, 8, 10, 10 | 0.0655 | 0.5089 | Sex x Stress | F (1, 34) = 0.02538 | 0.8744 | Unpaired t-test |
| | | | | | | Sex | F (1, 34) = 0.4281 | 0.5173 | |
| | | | | | | Stress | F (1, 34) = 8.551 | **0.0061** | |
| Figure 2i | 2-Way ANOVA | | 10, 8, 10, 10 | 0.2539 | 0.4340 | Sex x Stress | F (1, 34) = 3.256 | 0.0800 | None |
| | | | | | | Sex | F (1, 34) = 0.9128 | 0.3461 | |
| | | | | | | Stress | F (1, 34) = 0.008780 | 0.9259 | |
| Figure 3d | 2-Way RMANOVA | 11, 6 | 4, 2 | 0.2972 | N/A | Sex x CRF | F (1, 15) = 6.496 | **0.0223** | Paired t-test |
| | | | | | | Sex | F (1, 15) = 4.323 | 0.0552 | |
| | | | | | | CRF | F (1, 15) = 16.62 | **0.0010** | |
| Figure 3g | 3-Way RMANOVA | 12, 14, 11, 12 | 3, 3,3, 3 | 0.3658 | 0.0633 | CRF | F (1, 45) = 25.27 | **<0.0001** | Bonferroni |
| | | | | | | Sex | F (1, 45) = 0.2901 | 0.5928 | posthoc |
| | | | | | | Stress | F (1, 45) = 0.2617 | 0.6114 | |
| | | | | | | CRF x Sex | F (1, 45) = 3.842 | 0.0562 | |
| | | | | | | CRF x Stress | F (1, 45) = 2.447 | 0.1248 | |
| | | | | | | Sex x Stress | F (1, 45) = 5.192 | **0.0275** | |
| | | | | | | CRF x Sex x Stress | F (1, 45) = 5.696 | **0.0213** | |
| Figure 3i | 2-Way RMANOVA | 16, 14 | 3, 3 | **0.0004** | N/A | Sex x CRF | F (1, 28) = 14.97 | **0.0006** | Wilcoxon Signed- |
| | | | | | | Sex | F (1, 28) = 1.028 | 0.3193 | Rank Test (paired) |
| | | | | | | CRF | F (1, 28) = 13.44 | **0.0010** | |
| Figure 3k | 2-Way RMANOVA | 11, 13 | 3, 3 | 0.2633 | N/A | Sex x CRF | F (1, 22) = 12.49 | **0.0019** | Paired t-test |
| | | | | | | Sex | F (1, 22) = 0.001335 | 0.9712 | |
| | | | | | | CRF | F (1, 22) = 2.043 | 0.1669 | |

| | | | | | | | | | |
|---|---|---|---|---|---|---|---|---|---|
| Figure 4b | 2-Way ANOVA | 30, 33, 27, 32 | 5, 5, 4, 5 | **0.0000** | **<0.0001** | Sex x Stress | F (1, 118) = 6.635 | **0.0112** | Mann-Whitney |
| | | | | | | Sex | F (1, 118) = 0.0001346 | 0.9908 | |
| | | | | | | Stress | F (1, 118) = 4.705 | **0.0321** | |
| Figure 4c | 2-Way ANOVA | 30, 33, 27, 32 | 5, 5, 4, 5 | **0.0477** | **0.0041** | Sex x Stress | F (1, 118) = 0.3296 | 0.5670 | None |
| | | | | | | Sex | F (1, 118) = 0.2946 | 0.5883 | |
| | | | | | | Stress | F (1, 118) = 3.783 | 0.0542 | |
| Figure 4d | 2-Way ANOVA | 32, 35, 34, 34 | 5, 5, 4, 5 | **0.0147** | **<0.0001** | Sex x Stress | F (1, 131) = 9.885 | **0.0021** | Mann-Whitney |
| | | | | | | Sex | F (1, 131) = 0.004683 | 0.9455 | |
| | | | | | | Stress | F (1, 131) = 0.9300 | 0.3366 | |
| Figure 4e | 2-Way ANOVA | 27, 33,27, 32 | 5, 5, 4, 5 | 0.4357 | **0.0393** | Sex x Stress | F (1, 115) = 4.123 | **0.0446** | Mann-Whitney |
| | | | | | | Sex | F (1, 115) = 3.892 | 0.0509 | |
| | | | | | | Stress | F (1, 115) = 3.587 | 0.0608 | |
| Figure 4f | 2-Way ANOVA | 27, 33,27, 32 | 5, 5, 4, 5 | 0.2662 | 0.0882 | Sex x Stress | F (1, 115) = 1.802 | 0.1821 | None |
| | | | | | | Sex | F (1, 115) = 0.1059 | 0.7455 | |
| | | | | | | Stress | F (1, 115) = 2.334 | 0.1293 | |
| Figure 4g | 2-Way ANOVA | 29, 33, 28, 32 | 5, 5, 4, 5 | 0.0691 | **0.0002** | Sex x Stress | F (1, 118) = 0.001050 | 0.9742 | Mann-Whitney |
| | | | | | | Sex | F (1, 118) = 4.667 | **0.0328** | |
| | | | | | | Stress | F (1, 118) = 11.02 | **0.0012** | |
| Figure 4j-l | 3-Way ANOVA | 31, 34, 30, 32 | 5, 5, 4, 5 | **<0.0001** | **<0.0001** | CurrentInj | F (13, 1719) = 149.4 | **<0.0001** | |
| | | | | | | Sex | F (1, 1719) = 34.22 | **<0.0001** | |
| | | | | | | Stress | F (1, 1719) = 89.16 | **<0.0001** | |
| | | | | | | CurrentInj x Sex | F (13, 1719) = 0.7551 | 0.7086 | |
| | | | | | | CurrentInj x Stress | F (13, 1719) = 2.436 | **0.0029** | |
| | | | | | | Sex x Stress | F (1, 1719) = 31.50 | **<0.0001** | |
| | | | | | | CurrentInj x Sex x Stress | F (13, 1719) = 0.1435 | 0.9998 | |
| | 2-Way ANOVA - Males | | | | | Current Inj x Stress | F (13, 882) = 1.067 | 0.3844 | Mann-Whitney |
| | | | | | | Current Inj | F (13, 882) = 90.41 | **<0.0001** | |
| | | | | | | Stress | F (1, 882) = 10.24 | **0.0014** | |
| | 2-Way ANOVA - Females | | | | | Current Inj x Stress | F (13, 837) = 1.382 | 0.1615 | Mann-Whitney |
| | | | | | | Current Inj | F (13, 837) = 65.11 | **<0.0001** | |
| | | | | | | Stress | F (1, 837) = 86.39 | **<0.0001** | |
| Figure 4o-q | 3-Way ANOVA | 26, 29, 31, 30 | 4, 5, 4, 4 | **<0.0001** | **<0.0001** | CurrentInj | F (13, 1568) = 70.13 | **<0.0001** | |
| | | | | | | Sex | F (1, 1568) = 15.42 | **<0.0001** | |
| | | | | | | Stress | F (1, 1568) = 77.95 | **<0.0001** | |
| | | | | | | CurrentInj x Sex | F (13, 1568) = 0.7604 | 0.7030 | |
| | | | | | | CurrentInj x Stress | F (13, 1568) = 4.619 | **<0.0001** | |
| | | | | | | Sex x Stress | F (1, 1568) = 25.55 | **<0.0001** | |
| | | | | | | CurrentInj x Sex x Stress | F (13, 1568) = 1.296 | 0.2077 | |
| | 2-Way ANOVA - Males | | | | | Current Inj x Stress | F (13, 742) = 1.152 | **0.3109** | Mann-Whitney |
| | | | | | | Current Inj | F (13, 742) = 60.61 | **<0.0001** | |
| | | | | | | Stress | F (1, 742) = 15.08 | **0.0001** | |
| | 2-Way ANOVA - Females | | | | | Current Inj x Stress | F (13, 826) = 3.793 | **0.0000** | Mann-Whitney |
| | | | | | | Current Inj | F (13, 826) = 29.85 | **<0.0001** | |
| | | | | | | Stress | F (1, 826) = 68.06 | **<0.0001** | |
| Figure 4r,s | 3-Way ANOVA | 26, 29, 31, 30 | 4, 5, 4, 4 | **<0.0001** | **<0.0001** | CurrentInj | F (3, 436) = 2.485 | **0.0602** | |
| | | | | | | Sex | F (1, 436) = 12.41 | **0.0005** | |
| | | | | | | Stress | F (1, 436) = 65.52 | **<0.0001** | |
| | | | | | | CurrentInj x Sex | F (3, 436) = 0.04009 | 0.9893 | |
| | | | | | | CurrentInj x Stress | F (3, 436) = 0.8270 | 0.4795 | |
| | | | | | | Sex x Stress | F (1, 436) = 34.64 | **<0.0001** | |
| | | | | | | CurrentInj x Sex x Stress | F (3, 436) = 0.07611 | 0.9729 | |
| | 2-Way ANOVA - Males | | | | | Current Inj x Stress | F (3, 201) = 0.5231 | 0.6669 | None |
| | | | | | | Current Inj | F (3, 201) = 0.9726 | 0.4066 | |
| | | | | | | Stress | F (1, 201) = 2.345 | 0.1272 | |
| | 2-Way ANOVA - Females | | | | | Current Inj x Stress | F (3, 235) = 0.3747 | 0.7713 | Mann-Whitney |
| | | | | | | Current Inj | F (3, 235) = 1.588 | 0.1929 | |
| | | | | | | Stress | F (1, 235) = 102.9 | **<0.0001** | |

| | | | | | | | | | |
|---|---|---|---|---|---|---|---|---|---|
| Figure 5b, d | 3-Way ANOVA | 13, 14, 14, 12 | 3, 3, 3, 3 | **0.0078** | **<0.0001** | CurrentInj | F (3, 196) = 3.031 | **0.0305** | Mann Whitney |
| | | | | | | Sex | F (1, 196) = 0.02765 | 0.8681 | |
| | | | | | | Stress | F (1, 196) = 3.174 | 0.0764 | |
| | | | | | | CurrentInj x Sex | F (3, 196) = 0.01275 | 0.9980 | |
| | | | | | | CurrentInj x Stress | F (3, 196) = 0.003779 | 0.9997 | |
| | | | | | | Sex x Stress | F (1, 196) = 17.63 | **<0.0001** | |
| | | | | | | CurrentInj x Sex x Stress | F (3, 196) = 0.7622 | 0.5165 | |
| Figure 5c, e | 3-Way ANOVA | 13, 14, 14, 12 | 3, 3, 3, 3 | **<0.0001** | **<0.0001** | Rebox | F (3, 190) = 2.272 | 0.0815 | Mann Whitney |
| | | | | | | Sex | F (1, 190) = 3.458 | 0.0645 | |
| | | | | | | Stress | F (1, 190) = 2.465 | 0.1181 | |
| | | | | | | Rebox x Sex | F (3, 190) = 0.3967 | 0.7555 | |
| | | | | | | Rebox x Stress | F (3, 190) = 0.03949 | 0.9895 | |
| | | | | | | Sex x Stress | F (1, 190) = 2.790 | 0.0965 | |
| | | | | | | Rebox x Sex x Stress | F (3, 190) = 0.9828 | 0.4020 | |
| Figure 4f | 3-Way RMANOVA | 12, 14, 11, 12 | 3, 3,3, 3 | **<0.0001** | **<0.0001** | 4AP | F (1, 47) = 33.10 | **<0.0001** | Wilcoxon Ranked |
| | | | | | | Sex | F (1, 47) = 0.1191 | 0.7316 | |
| | | | | | | Stress | F (1, 47) = 0.3480 | 0.5581 | |
| | | | | | | 4AP x Sex | F (1, 47) = 1.046 | 0.3116 | |
| | | | | | | 4AP x Stress | F (1, 47) = 0.2023 | 0.6550 | |
| | | | | | | Sex x Stress | F (1, 47) = 9.325 | **0.0037** | |
| | | | | | | 4AP x Sex x Stress | F (1, 47) = 7.362 | **0.0093** | |

| | | | | | | | | | |
|---|---|---|---|---|---|---|---|---|---|
| Figure 6b | 3-Way RMANOVA | | 8, 10, 7, 8, 8, 8, 6, 7 | 0.3733 | 0.7357 | Rebox | F (1, 54) = 0.4793 | **0.4900** | Bonferroni |
| | | | | | | Sex | F (1, 54) = 2.393 | 0.1300 | Posthoc |
| | | | | | | Stress | F (1, 54) = 12.37 | **<0.001** | |
| | | | | | | Rebox x Sex | F (1, 54) = 6.255 | **0.0200** | |
| | | | | | | Rebox x Stress | F (1, 54) = 26.75 | **<0.001** | |
| | | | | | | Sex x Stress | F (1, 54) = 2.095 | 0.1500 | |
| | | | | | | Rebox x Sex x Stress | F (1, 54) = 4.698 | **0.0300** | |
| Figure 6d | 3-Way RMANOVA | | 8 ,6, 9, 8, 8, 8, 6, 7 | 0.3047 | **0.0486** | Rebox | F (1, 52) = 7.846 | **0.0071** | |
| | | | | | | Sex | F (1, 52) = 1.917 | 0.1721 | |
| | | | | | | Stress | F (1, 52) = 0.8031 | 0.3743 | |
| | | | | | | Rebox x Sex | F (1, 52) = 0.2515 | 0.6182 | |
| | | | | | | Rebox x Stress | F (1, 52) = 0.2056 | 0.6521 | |
| | | | | | | Sex x Stress | F (1, 52) = 0.3966 | 0.5316 | |
| | | | | | | Rebox x Sex x Stress | F (1, 52) = 1.029 | 0.3151 | |
| | 2-Way RMANOVA | | 16, 14, 15, 15 | | | Rebox x Stress | F (1, 56) = 0.2783 | 0.5999 | Mann-Whitney |
| | | | | | | Rebox | F (1, 56) = 7.037 | **0.0104** | |
| | | | | | | Stress | F (1, 56) = 0.5078 | 0.4791 | |
| Figure 6f | 3-Way RMANOVA | | 16, 14, 15, 15 | 0.2688 | 0.7984 | Rebox | F (1, 52) = 0.1815 | 0.6718 | None |
| | | | | | | Sex | F (1, 52) = 0.9107 | 0.3444 | |
| | | | | | | Stress | F (1, 52) = 0.1041 | 0.7482 | |
| | | | | | | Rebox x Sex | F (1, 52) = 0.4278 | 0.5160 | |
| | | | | | | Rebox x Stress | F (1, 52) = 2.345 | 0.1317 | |
| | | | | | | Sex x Stress | F (1, 52) = 0.02560 | 0.8735 | |
| | | | | | | Rebox x Sex x Stress | F (1, 52) = 0.2172 | 0.6431 | |

| | | | | | | | | | |
|---|---|---|---|---|---|---|---|---|---|
| Supplemental Figure 1b | 2-Way ANOVA | | 14, 10, 13, 13 | 0.3433 | 0.9834 | Sex x Stress | F (1, 46) = 0.3255 | 0.5711 | None |
| | | | | | | Sex | F (1, 46) = 0.001033 | 0.9745 | |
| | | | | | | Stress | F (1, 46) = 1.017 | 0.3185 | |
| Supplemental Figure 1c | 2-Way ANOVA | | 14, 10, 13, 13 | 0.4046 | 0.9695 | Sex x Stress | F (1, 45) = 6.103 | **0.0174** | Unpaired t-test |
| | | | | | | Sex | F (1, 45) = 0.07416 | 0.7866 | |
| | | | | | | Stress | F (1, 45) = 0.009729 | 0.9219 | |
| Supplemental Figure 1d | 2-Way ANOVA | | 14, 10, 13, 13 | 0.1592 | 0.9671 | Sex x Stress | F (1, 45) = 0.1704 | 0.6817 | None |
| | | | | | | Sex | F (1, 45) = 2.659 | 0.1100 | |
| | | | | | | Stress | F (1, 45) = 3.156 | 0.0824 | |
| Supplemental Figure 1f, g | 3-Way RMANOVA | | 9, 13, 12, 9 | 0.04 | 0.06 | Trial | F (4.167, 162.5) = 25.76 | **<0.001** | None |
| | | | | | | Sex | F (1, 39) = 0.07930 | 0.7800 | |
| | | | | | | Stress | F (1, 39) = 0.07764 | 0.7800 | |
| | | | | | | Trial x Sex | F (5, 195) = 1.465 | 0.2000 | |
| | | | | | | Trial x Stress | F (5, 195) = 0.7796 | 0.5700 | |
| | | | | | | Sex x Stress | F (1, 39) = 0.4192 | 0.5200 | |
| | | | | | | Trial x Sex x Stress | F (5, 195) = 1.317 | 0.2600 | |

| | | | | | | | | | |
|---|---|---|---|---|---|---|---|---|---|
| Supplemental Figure 3a | 2-Way RMANOVA | 8, 9 | 2, 3 | **0.0137** | 0.0504 | Sex x CRF | F (1, 15) = 0.3885 | 0.5424 | Wilcoxon Signed-Rank Test (paired) |
| | | | | | | Sex | F (1, 15) = 0.9781 | 0.3384 | |
| | | | | | | CRF | F (1, 15) = 11.09 | **0.0046** | |
| Supplemental Figure 3b | 2-Way RMANOVA | 6, 10 | 2, 3 | 0.0580 | **0.0133** | Sex x CRF | F (1, 28) = 0.06534 | 0.8001 | None |
| | | | | | | Sex | F (1, 28) = 0.07220 | 0.7901 | |
| | | | | | | CRF | F (1, 28) = 0.0001346 | 0.9908 | |

| | | | | | | | | | |
|---|---|---|---|---|---|---|---|---|---|
| Supplemental Figure 4a, b | 3-Way RMANOVA | 9, 11 | 3, 3 | **<0.0001** | 0.5299 | CurrentInj | F (1.523, 27.42) = 131.0 | **<0.0001** | |
| | | | | | | Sex | F (1, 18) = 1.449 | 0.2442 | |
| | | | | | | CRF | F (1.000, 18.00) = 7.872 | **0.0117** | |
| | | | | | | CurrentInj x Sex | F (13, 234) = 0.5085 | 0.9183 | |
| | | | | | | CurrentInj x CRF | F (2.570, 46.27) = 1.768 | 0.1733 | |
| | | | | | | Sex x CRF | F (1, 18) = 1.092 | 0.3098 | |
| | | | | | | CurrentInj x Sex x CRF | F (13, 234) = 0.4065 | 0.9667 | |
| | 2-Way RMANOVA - Males | | | | | CurrentInj | F (1.139, 9.110) = 43.20 | **<0.0001** | None |
| | | | | | | CRF | F (1.000, 8.000) = 3.961 | 0.0817 | |
| | | | | | | CurrentInj x CRF | F (1.477, 11.82) = 1.100 | 0.3446 | |
| | 2-Way RMANOVA - Females | | | | | CurrentInj | F (1.467, 14.67) = 114.0 | **<0.0001** | None |
| | | | | | | CRF | F (1.000, 10.00) = 3.925 | 0.0757 | |
| | | | | | | CurrentInj x CRF | F (1.572, 15.72) = 0.8425 | 0.4233 | |
| Supplemental Figure 4c, d | 3-Way RMANOVA | 10, 11 | 3, 3 | **<0.0001** | **<0.0001** | CurrentInj | F (1.699, 32.28) = 213.0 | **<0.0001** | |
| | | | | | | Sex | F (1, 19) = 0.7295 | 0.4037 | |
| | | | | | | CRF | F (1.000, 19.00) = 0.08726 | 0.7709 | |
| | | | | | | CurrentInj x Sex | F (13, 247) = 0.4754 | 0.9370 | |
| | | | | | | CurrentInj x CRF | F (2.722, 51.72) = 0.1369 | 0.9243 | |
| | | | | | | Sex x CRF | F (1, 19) = 0.7571 | 0.3951 | |
| | | | | | | CurrentInj x Sex x CRF | F (13, 247) = 0.3896 | 0.9724 | |
| | 2-Way RMANOVA - Males | | | | | CurrentInj | F (2.635, 23.72) = 1055 | **<0.0001** | None |
| | | | | | | CRF | F (1.000, 9.000) = 0.7747 | 0.4016 | |
| | | | | | | CurrentInj x CRF | F (4.012, 36.11) = 0.6304 | 0.6445 | |
| | 2-Way RMANOVA - Females | | | | | CurrentInj | F (13, 130) = 60.44 | **<0.0001** | None |
| | | | | | | CRF | F (1, 10) = 0.4154 | 0.5338 | |
| | | | | | | CurrentInj x CRF | F (13, 130) = 0.2185 | 0.9982 | |
| Supplemental Figure 4e, f | 3-Way RMANOVA | 12, 13 | 3, 3 | **<0.0001** | **<0.0001** | CurrentInj | F (2.452, 56.40) = 235.0 | **<0.0001** | |
| | | | | | | Sex | F (1, 23) = 0.9555 | 0.3385 | |
| | | | | | | CRF | F (1.000, 23.00) = 0.1649 | 0.6885 | |
| | | | | | | CurrentInj x Sex | F (13, 299) = 0.7703 | 0.6915 | |
| | | | | | | CurrentInj x CRF | F (2.372, 54.55) = 0.3486 | 0.7429 | |
| | | | | | | Sex x CRF | F (1, 23) = 4.380 | **0.0476** | |
| | | | | | | CurrentInj x Sex x CRF | F (13, 299) = 0.8842 | 0.5703 | |
| | 2-Way RMANOVA - Males | | | | | CurrentInj | F (2.087, 22.95) = 120.7 | **<0.0001** | None |
| | | | | | | CRF | F (1.000, 11.00) = 1.093 | 0.3183 | |
| | | | | | | CurrentInj x CRF | F (1.522, 16.74) = 0.4455 | 0.5952 | |
| | 2-Way RMANOVA - Females | | | | | CurrentInj | F (1.585, 19.02) = 118.2 | **0.0001** | None |
| | | | | | | CRF | F (1.000, 12.00) = 4.228 | 0.0622 | |
| | | | | | | CurrentInj x CRF | F (2.026, 24.31) = 1.231 | 0.3099 | |
| Supplemental Figure 4g, h | 3-Way RMANOVA | 11, 13 | 3, 3 | **<0.0001** | **<0.0001** | CurrentInj | F (2.539, 55.86) = 306.9 | **<0.0001** | |
| | | | | | | Sex | F (1, 22) = 0.1754 | 0.6794 | |
| | | | | | | CRF | F (1.000, 22.00) = 0.6237 | 0.4381 | |
| | | | | | | CurrentInj x Sex | F (13, 286) = 0.1954 | 0.9991 | |
| | | | | | | CurrentInj x CRF | F (2.443, 53.74) = 1.072 | 0.3597 | |
| | | | | | | Sex x CRF | F (1, 22) = 9.758 | **0.0049** | |
| | | | | | | CurrentInj x Sex x CRF | F (13, 286) = 5.031 | **<0.0001** | |
| | 2-Way RMANOVA - Males | | | | | CurrentInj | F (2.290, 22.90) = 230.2 | **<0.0001** | Wilcoxon Signed-Rank Test (paired) |
| | | | | | | CRF | F (1.000, 10.00) = 9.231 | **0.0125** | |
| | | | | | | CurrentInj x CRF | F (2.583, 25.83) = 4.546 | **0.0140** | |
| | 2-Way RMANOVA - Females | | | | | CurrentInj | F (1.663, 19.95) = 129.7 | **<0.0001** | None |
| | | | | | | CRF | F (1.000, 12.00) = 2.486 | 0.1408 | |
| | | | | | | CurrentInj x CRF | F (1.912, 22.94) = 1.489 | 0.2464 | |

| | | | | | | | | | |
|---|---|---|---|---|---|---|---|---|---|
| Supplemental Figure 5b | 2-Way ANOVA | 11, 14, 15, 17 | 3, 3, 4, 4 | 0.032455 | 0.140662 | Sex x Stress | F (1, 53) = 1.657 | 0.2037 | None |
| | | | | | | Sex | F (1, 53) = 0.07577 | 0.7842 | |
| | | | | | | Stress | F (1, 53) = 3.070 | 0.0855 | |
| Supplemental Figure 5c | 2-Way ANOVA | 11, 14, 15, 17 | 3, 3, 4, 4 | **0.0178** | **0.0001** | Sex x Stress | F (1, 53) = 2.295 | 0.1357 | None |
| | | | | | | Sex | F (1, 53) = 1.371 | 0.2468 | |
| | | | | | | Stress | F (1, 53) = 0.1946 | 0.6609 | |
| Supplemental Figure 5e | 2-Way ANOVA | 9, 10, 9, 16 | 3, 3, 4, 4 | 0.4435 | **0.0000** | Sex x Stress | F (1, 40) = 0.009521 | 0.9228 | None |
| | | | | | | Sex | F (1, 40) = 0.2566 | 0.6152 | |
| | | | | | | Stress | F (1, 40) = 0.7747 | 0.3840 | |
| Supplemental Figure 5f | 2-Way ANOVA | 9, 10, 9, 16 | 3, 3, 4, 4 | **0.0006** | **0.0124** | Sex x Stress | F (1, 40) = 1.456 | 0.2347 | None |
| | | | | | | Sex | F (1, 40) = 2.446 | 0.1257 | |
| | | | | | | Stress | F (1, 40) = 0.08228 | 0.7757 | |
| Supplemental Figure 6b | 2-Way ANOVA | 12, 13, 11, 12 | 3, 3, 3, 3 | 0.2704 | **0.0359** | Sex x Stress | F (1, 45) = 0.9425 | 0.3368 | None |
| | | | | | | Sex | F (1, 45) = 0.1102 | 0.7415 | |
| | | | | | | Stress | F (1, 45) = 2.494 | 0.1213 | |
| Supplemental Figure 6g,h | 3-Way ANOVA | 9, 10, 11, 11 | 3, 3, 3, 3 | **<0.0001** | **<0.0001** | CurrentInj | F (13, 518) = 119.2 | **<0.0001** | |
| | | | | | | Sex | F (1, 518) = 5.064 | **0.0248** | |
| | | | | | | Stress | F (1, 518) = 6.361 | **0.0120** | |
| | | | | | | CurrentInj x Sex | F (13, 518) = 0.1576 | 0.9997 | |
| | | | | | | CurrentInj x Stress | F (13, 518) = 0.1954 | 0.9991 | |
| | | | | | | Sex x Stress | F (1, 518) = 0.09221 | 0.7615 | |
| | | | | | | CurrentInj x Sex x Stress | F (13, 518) = 0.05794 | 1.0000 | |
| | 2-Way ANOVA - Males | | | | | Current Inj x Stress | F (13, 238) = 0.1229 | 1.0000 | None |
| | | | | | | Current Inj | F (13, 238) = 56.41 | **<0.0001** | |
| | | | | | | Stress | F (1, 238) = 2.241 | 0.1400 | |
| | 2-Way ANOVA - Females | | | | | Current Inj x Stress | F (13, 280) = 0.1303 | 1.0000 | Mann-Whitney |
| | | | | | | Current Inj | F (13, 280) = 63.25 | **<0.0001** | |
| | | | | | | Stress | F (1, 280) = 4.396 | **0.0400** | |

| | | | | | | | | | |
|---|---|---|---|---|---|---|---|---|---|
| Supplemental Figure 7b | 2-Way ANOVA | 27, 20, 31, 31 | 4, 5, 4, 4 | **0.0006** | **<0.0001** | Sex x Stress | F (1, 105) = 0.4216 | 0.5176 | Mann-Whitney |
| | | | | | | Sex | F (1, 105) = 5.356 | **0.0226** | |
| | | | | | | Stress | F (1, 105) = 13.62 | **0.0004** | |
| Supplemental Figure 7c | 2-Way ANOVA | 28, 21, 31, 31 | 4, 5, 4, 4 | 0.2439 | **0.0002** | Sex x Stress | F (1, 107) = 0.0002082 | 0.9885 | None |
| | | | | | | Sex | F (1, 107) = 6.634 | **0.0114** | |
| | | | | | | Stress | F (1, 107) = 0.4148 | 0.5209 | |
| Supplemental Figure 7d | 2-Way ANOVA | 34, 30, 37, 32 | 4, 5, 4, 4 | **0.0000** | **<0.000001** | Sex x Stress | F (1, 129) = 7.031 | **0.0090** | Mann-Whitney |
| | | | | | | Sex | F (1, 129) = 0.6555 | 0.4196 | |
| | | | | | | Stress | F (1, 129) = 0.03772 | 0.8463 | |
| Supplemental Figure 7e | 2-Way ANOVA | 26, 20, 31, 31 | 4, 5, 4, 4 | **0.0074** | 0.2428 | Sex x Stress | F (1, 105) = 0.3991 | 0.5289 | Mann-Whitney |
| | | | | | | Sex | F (1, 105) = 4.045 | **0.0469** | |
| | | | | | | Stress | F (1, 105) = 5.188 | **0.0248** | |
| Supplemental Figure 7f | 2-Way ANOVA | 26, 20, 31, 31 | 4, 5, 4, 4 | 0.0815 | 0.4407 | Sex x Stress | F (1, 105) = 0.07270 | 0.7880 | None |
| | | | | | | Sex | F (1, 105) = 2.024 | 0.1578 | |
| | | | | | | Stress | F (1, 105) = 0.02628 | 0.8715 | |
| Supplemental Figure 7g | 2-Way ANOVA | 26, 20, 31, 31 | 4, 5, 4, 4 | **0.0159** | **0.0000** | Sex x Stress | F (1, 105) = 0.001744 | 0.9668 | None |
| | | | | | | Sex | F (1, 105) = 0.0004339 | 0.9834 | |
| | | | | | | Stress | F (1, 105) = 2.136 | 0.1469 | |
| Supplemental Figure 7j-l | 3-Way ANOVA | 33, 30, 37, 32 | 4, 5, 4, 4 | **<0.0001** | **<0.0001** | CurrentInj | F (13, 1792) = 502.7 | **<0.0001** | |
| | | | | | | Sex | F (1, 1792) = 11.10 | **0.0009** | |
| | | | | | | Stress | F (1, 1792) = 59.93 | **<0.0001** | |
| | | | | | | CurrentInj x Sex | F (13, 1792) = 1.789 | **0.0396** | |
| | | | | | | CurrentInj x Stress | F (13, 1792) = 0.5010 | 0.9251 | |
| | | | | | | Sex x Stress | F (1, 1792) = 24.83 | **<0.0001** | |
| | | | | | | CurrentInj x Sex x Stress | F (13, 1792) = 0.2220 | 0.9983 | |
| | 2-Way ANOVA - Males | | | | | Current Inj x Stress | F (13, 854) = 0.5074 | 0.9210 | None |
| | | | | | | CurrentInj | F (13, 854) = 237.9 | **<0.0001** | |
| | | | | | | Stress | F (1, 854) = 3.263 | 0.0712 | |
| | 2-Way ANOVA - Females | | | | | Current Inj x Stress | F (13, 938) = 0.1538 | 0.9998 | Mann-Whitney |
| | | | | | | CurrentInj | F (13, 938) = 265.9 | **<0.0001** | |
| | | | | | | Stress | F (1, 938) = 94.76 | **<0.0001** | |
| Supplemental Figure 7o-q | 3-Way ANOVA | 32, 33, 30, 33 | 4, 5, 4, 4 | **<0.0001** | **<0.0001** | CurrentInj | F (13, 1736) = 308.4 | **<0.0001** | |
| | | | | | | Sex | F (1, 1736) = 54.96 | **<0.0001** | |
| | | | | | | Stress | F (1, 1736) = 19.41 | **<0.0001** | |
| | | | | | | CurrentInj x Sex | F (13, 1736) = 3.098 | **0.0001** | |
| | | | | | | CurrentInj x Stress | F (13, 1736) = 1.094 | 0.3593 | |
| | | | | | | Sex x Stress | F (1, 1736) = 12.34 | **0.0005** | |
| | | | | | | CurrentInj x Sex x Stress | F (13, 1736) = 1.207 | 0.2671 | |
| | 2-Way ANOVA - Males | | | | | Current Inj x Stress | F (13, 882) = 0.05451 | 1.0000 | None |
| | | | | | | CurrentInj | F (13, 882) = 130.9 | **<0.0001** | |
| | | | | | | Stress | F (1, 882) = 0.2802 | 0.5967 | |
| | 2-Way ANOVA - Females | | | | | Current Inj x Stress | F (13, 854) = 4.070 | **<0.0001** | Mann-Whitney |
| | | | | | | CurrentInj | F (13, 854) = 229.0 | **<0.0001** | |
| | | | | | | Stress | F (1, 854) = 57.37 | **<0.0001** | |
| Supplemental Figure 7r,s | 3-Way ANOVA | 32, 33, 30, 33 | 4, 5, 4, 4 | **<0.0001** | **<0.0001** | CurrentInj | F (3, 492) = 3.297 | **0.0203** | |
| | | | | | | Sex | F (1, 492) = 66.46 | **<0.0001** | |
| | | | | | | Stress | F (1, 492) = 13.08 | **0.0003** | |
| | | | | | | CurrentInj x Sex | F (3, 492) = 0.3477 | 0.7908 | |
| | | | | | | CurrentInj x Stress | F (3, 492) = 0.02476 | 0.9947 | |
| | | | | | | Sex x Stress | F (1, 492) = 67.66 | **<0.0001** | |
| | | | | | | CurrentInj x Sex x Stress | F (3, 492) = 0.2168 | 0.8847 | |
| | 2-Way ANOVA - Males | | | | | Current Inj x Stress | F (3, 248) = 0.1381 | 0.9372 | Mann-Whitney |
| | | | | | | CurrentInj | F (3, 248) = 1.326 | 0.2664 | |
| | | | | | | Stress | F (1, 248) = 18.40 | **<0.0001** | |
| | 2-Way ANOVA - Females | | | | | Current Inj x Stress | F (3, 244) = 0.1127 | 0.9526 | Mann-Whitney |
| | | | | | | CurrentInj | F (3, 244) = 2.004 | 0.1141 | |
| | | | | | | Stress | F (1, 244) = 48.80 | **<0.0001** | |
